# Identification of EcpK, a bacterial tyrosine pseudokinase important for exopolysaccharide biosynthesis in *Myxococcus xanthus*

**DOI:** 10.1101/2024.11.26.625375

**Authors:** Luca Blöcher, Johannes Schwabe, Timo Glatter, Lotte Søgaard-Andersen

## Abstract

Bacteria synthesize and export chemically diverse polysaccharides that function in many physiological processes and are widely used in industrial applications. In the ubiquitous Wzx/Wzy-dependent biosynthetic pathways for these polysaccharides, the polysaccharide co-polymerase (PCP) facilitates the polymerization of repeat units in the periplasm, and in Gram-negative bacteria, also polysaccharide translocation across the outer membrane. These PCPs are integral inner membrane proteins with extended periplasmic domains, and functionally depend on alternating between different oligomeric states. The oligomeric state, in turn, is determined by a cognate cytoplasmic bacterial tyrosine kinase (BYK), which is either part of the PCP or a stand-alone protein. Interestingly, BYK-like proteins, which lack key catalytic residues and/or the phosphorylated Tyr residues, have been described. In *Myxococcus xanthu*s, the exopolysaccharide (EPS) is synthesized and exported *via* the Wzx/Wzy-dependent EPS pathway in which EpsV serves as the PCP. Here, we confirm that EpsV lacks the BYK domain. Using phylogenomics, experiments and computational structural biology, we identify EcpK as important for EPS biosynthesis and show that it structurally resembles canonical BYKs but lacks residues important for catalysis and Tyr phosphorylation. Using proteomic analyses, two-hybrid assays and structural modeling, we demonstrate that EcpK directly interacts with EpsV. Based on these findings, we suggest that EcpK is a BY pseudokinase and functions as a scaffold, which by direct protein-protein interactions, rather than by Tyr phosphorylation, facilitates EpsV function. EcpK and EpsV homologs are present in other bacteria, suggesting a broad conservation of this mechanism.

**Importance:** Bacteria produce a variety of polysaccharides with important biological functions. In Wzx/Wzy-dependent pathways for the biosynthesis of secreted and capsular polysaccharides in Gram-negative bacteria, the polysaccharide co-polymerase (PCP) is a key protein that facilitates repeat unit polymerization and polysaccharide translocation across the outer membrane. PCP function depends on assembly/disassembly cycles that are determined by the phosphorylation/dephosphorylation cycles of an associated bacterial tyrosine kinase (BYK). Here, we identify the BY pseudokinase EcpK as essential for exopolysaccharide biosynthesis in *Myxococcus xanthus*. Based on experiments and computational structural biology, we suggest that EcpK is a scaffold protein, guiding the assembly/disassembly cycles of the partner PCP *via* binding/unbinding cycles independently of Tyr phosphorylation/dephosphorylation cycles. We suggest that this novel mechanism is broadly conserved.

## Introduction

Bacteria produce and export a range of different polysaccharides essential for various biological functions, including biofilm formation, virulence, adhesion, motility, host-microbe interactions, while also providing protection against phage infection and external stresses (1-3). Many of these polysaccharides also have applications in the food, biomedical, and pharmaceutical industries (4).

Bacterial polysaccharide biosynthesis and export occur *via* three common pathways, i.e., the Wzx/Wzy-dependent, the ABC transporter-dependent, and the synthase-dependent pathways (5-7). In the ubiquitous Wzx/Wzy-dependent pathway, synthesis begins at the cytoplasmic leaflet of the inner membrane (IM) by a phosphoglycosyltransferase (PGT), which transfers a sugar-1 phosphate from a nucleotide-sugar precursor to undecaprenyl phosphate (Und-P) to form an Und-PP-sugar molecule (5). Next, glycosyltransferases (GTs) add monosaccharides to synthesize the repeat unit, which is then translocated to the outer leaflet of the IM periplasm by the Wzx flippase (5). Subsequently, the Wzy polymerase polymerizes the repeat units (5). For the promotion of polymerization, an additional integral IM protein, the polysaccharide co-polymerase (PCP), is required to work in conjunction with the Wzy polymerase (8-14). Although the mechanism by which the PCP controls polymerization is not fully understood, it likely involves direct interaction with the Wzy polymerase (8, 13-19). Finally, in Gram-negative bacterial pathways for capsular and secreted polysaccharides, polysaccharides are translocated across the outer membrane (OM) by either an integral OM polysaccharide export (OPX) protein or a composite translocon comprising an integral OM β-barrel protein and a periplasmic OPX protein (5, 20, 21) in a process that depends on the PCP and involves direct contact between the PCP and the OPX protein (22-24). Thus, in Wzx/Wzy-dependent pathways for capsular and secreted polysaccharide biosynthesis in Gram-negative bacteria, the PCP is a key protein acting at the nexus between repeat unit polymerization and polysaccharide translocation across the OM.

PCPs in Wzx/Wzy-dependent pathways are integral IM proteins and are classified into two families (25). PCP-1 proteins function in Wzx/Wzy-dependent pathways for enterobacterial common antigen and the O-antigen of LPS (25-27). These proteins comprise two transmembrane helices (TMH) and a large mostly α-helical periplasmic domain, and assemble to form octameric complexes, with the 16 TMHs creating a cage-like structure in the IM, and the periplasmic domains forming a tapered cavity closed at the top (19, 28-31). PCP-2 proteins function in Wzx/Wzy-dependent biosynthetic pathways for capsular and secreted polysaccharides (5, 25). PCP-2 proteins also comprise two TMHs and a large, mostly α-helical periplasmic domain (25, 32). As a distinguishing characteristic, they also incorporate a cytoplasmic bacterial tyrosine kinase (BYK), which is crucial for their function (10, 33-40). Most Wzx/Wzy-dependent pathways in Gram-negative bacteria incorporate a PCP-2a protein in which the PCP part is fused to the BYK domain in a single polypeptide (5, 25, 41-43). By contrast, bipartite PCP-2b proteins function together with a separate, stand-alone BYK, and are mostly found in Gram-positive bacteria but also in some Gram-negative bacteria (5, 25, 41-43). Unlike tyrosine kinases in eukaryotes, BYKs have a Rossmann-like fold, structurally resemble P-loop ATPases, and contain at their C-terminus a flexible loop with multiple Tyr-residues, i.e. the Tyr-rich tail, which serve as substrates for phosphorylation (10, 32, 44-52).

The structure and function of PCP-2a proteins are exemplified by Wzc of *Escherichia coli* (Wzc*^E.coli^*), which is important for the polymerization and translocation across the OM of capsular polysaccharide (10, 22-24, 32, 53-55). Wzc*^E.coli^* forms octamers under conditions in which no or only a few Tyr-residues in the Tyr-rich tails are phosphorylated (32, 49). In this octamer, the cytoplasmic BYK domains are arranged in a ring-like structure in which each protomer’s Tyr-rich tail reaches into the neighboring protomer’s active site, facilitating transphosphorylation (32, 49). Moreover, the 16 TMHs create a cage-like structure in the IM, and the eight periplasmic domains form a tapered cavity open at the top (32), likely involved in interaction with the OPX protein Wza*^E.coli^* (22-24). Increased phosphorylation of the Tyr-residues destabilize the octamer to the point of dissociation (32, 49). Tyr phosphorylation is counteracted by the Wzb*^E.coli^* phospho-tyrosine phosphatase, thereby restoring octamer formation (10, 33, 40, 56-58).

The CapAB proteins of *Staphylococcus aureus* (CapAB*^S. aureus^*^)^, which are essential for capsule biosynthesis, exemplify the structure and function of bipartite PCP-2b proteins with their associated BYKs (34, 37, 51, 59). CapB*^S. aureus^* BYK activity is activated by direct interaction with the short C-terminal cytoplasmic extension of its PCP-2b transmembrane/extracytoplasmic partner protein, CapA*^S. aureus^* (34). Non-phosphorylated CapB*^S. aureus^* fused to this extension also assembles to form an octameric ring in which the Tyr-rich tail from one protomer also extends into the active site of a neighboring protomer, thereby facilitating transphosphorylation (51). Also in this system, transphosphorylation of these Tyr-residues causes the dissociation of the octamer (51). This process is reversed by the functionally important CapC*^S. aureus^* phospho-tyrosine phosphatase, which catalyzes the dephosphorylation of CapB*^S. aureus^* Tyr-residues (37, 51, 60, 61). Thus, in the current model for pathways incorporating a PCP-2, the phosphorylation/dephosphorylation cycles that facilitate the alternation between oligomeric states are essential for PCP-2 function (19, 32, 48, 49, 56, 57, 60). Nevertheless, sequence based analyses and experimental work on the HfsAB proteins in *Caulobacter crescentus* have identified non-canonical PCP-2a proteins and stand-alone BYK domains that lack residues essential for catalytic activity (25, 62).

In *Myxococcus xanthus,* the exopolysaccharide (EPS) is crucial for several biological traits, including type IV pili (T4P)-dependent motility, adhesion, development and biofilm formation (63). EPS is synthesized and exported by the Wzx/Wzy-dependent EPS pathway (20, 21, 64-66). In this pathway, repeat unit biosynthesis is initiated by the PGT EpsZ, which transfers galactose-1-P to Und-P (64, 66). Subsequently, several GTs add monosaccharides to finalize the EPS repeat unit, which is translocated into the periplasm by the Wzx_EPS_ flippase. The Wzy_EPS_ polymerase builds the final EPS by polymerizing the repeat units (21, 64-66). EPS is translocated across the OM by a bipartite translocon comprising the periplasmic ^D1D2^OPX protein EpsY and the integral OM β-barrel protein EpsX (20, 21). EpsV is the PCP-2 protein of this pathway; however, it does not contain a BYK domain (20, 21, 25, 64, 66). MXAN_7447 was proposed to be its stand-alone BYK partner (66).

Here, using computational analyses and experiments, we characterize MXAN_7447 as a BY pseudokinase important for EPS biosynthesis. We further demonstrate that MXAN_7447 (henceforth named <u>E</u>psV-<u>c</u>oupled BY <u>p</u>seudo<u>k</u>inase, EcpK) interacts directly with the PCP-2 of the EPS biosynthetic pathway, EpsV, which lacks a BYK domain. We speculate that EcpK functions as a scaffold to facilitate EpsV assembly/disassembly *via* direct interactions. We also identified orthologs of EcpK and EpsV in other Myxobacteria. These findings together with previous sequence-based analyses (25) and experimental analyses of the HfsAB proteins in *C. crescentus* (62) suggest that PCP-2 proteins that function with a BY pseudokinase are widespread. Our findings provide evidence for an alternative mechanism of PCP-2 function involving a BY pseudokinase, thus diverging from the canonical BYK-dependent phosphorylation mechanism.

## Results

### Myxobacterial gene clusters for EPS biosynthesis encode a BY pseudokinase

The genes encoding the EPS biosynthetic pathway are present in two gene clusters separated by 18 genes and are conserved in related Myxobacteria (21, 64-66) (Fig. S1A and B). In agreement with previous sequence-based analyses, *epsV* (*MXAN_7421*) and its orthologs encode PCP-2 proteins lacking the BYK domain (Fig. S1B and C) (20, 21, 25, 66). Consistently, previous analyses based on AlphaFold2 structural models documented that the EpsV monomer has a structure similar to that of a Wzc*^E. coli^* protomer in the solved octameric structure of Wzc*^E. coli^*, except that EpsV lacks the cytoplasmic BYK domain and has a larger periplasmic domain (20, 21, 32).

Typically, for bipartite PCP-2b proteins, the PCP-2b/BYK pair is encoded by neighboring genes (43); however, none of the genes adjacent to *epsV* encode a BYK (Fig. S1A and B). Previously, it was proposed that EcpK (MXAN_7447), which was suggested to be important for EPS biosynthesis, is the stand-alone BYK partner of EpsV (66). *epsV* and *ecpK* orthologs largely co-occur in the two *eps* gene clusters (Fig. S1A and B). To explore a possible function of EcpK in EPS biosynthesis, we first generated an AlphaFold2 (67) structural model (Fig. 1A and Fig. S2). In this high-confidence model, EcpK adopts a Rossmann fold-like structure consisting of a central five-stranded β-sheet surrounded by five α-helices, with an unstructured N-terminal region and an additional α-helix at the C-terminal end (Fig. 1A). To identify structural homologs of EcpK, we performed a Foldseek (68) search in the PDB database. We identified the BYK domain of full-length Wzc*^E.coli^* in the octameric structure (32), the BYK domains of the Wzc-like monomeric Etk*^E.coli^* (50) and dimeric VpsO*^Vibrio cholerae^* (48), as well as the BYK domain of the CapAB*^S. aureus^* octamer (51), as structural homologs of EcpK (Fig. 1B and C). EcpK’s central five-stranded β-sheet and the surrounding five α-helices aligned well with the corresponding structures in the solved structures of these four BYKs, while the unstructured N-terminus and the C-terminal α-helix of EcpK did not (Fig. 1B and C).

**Figure 1.**
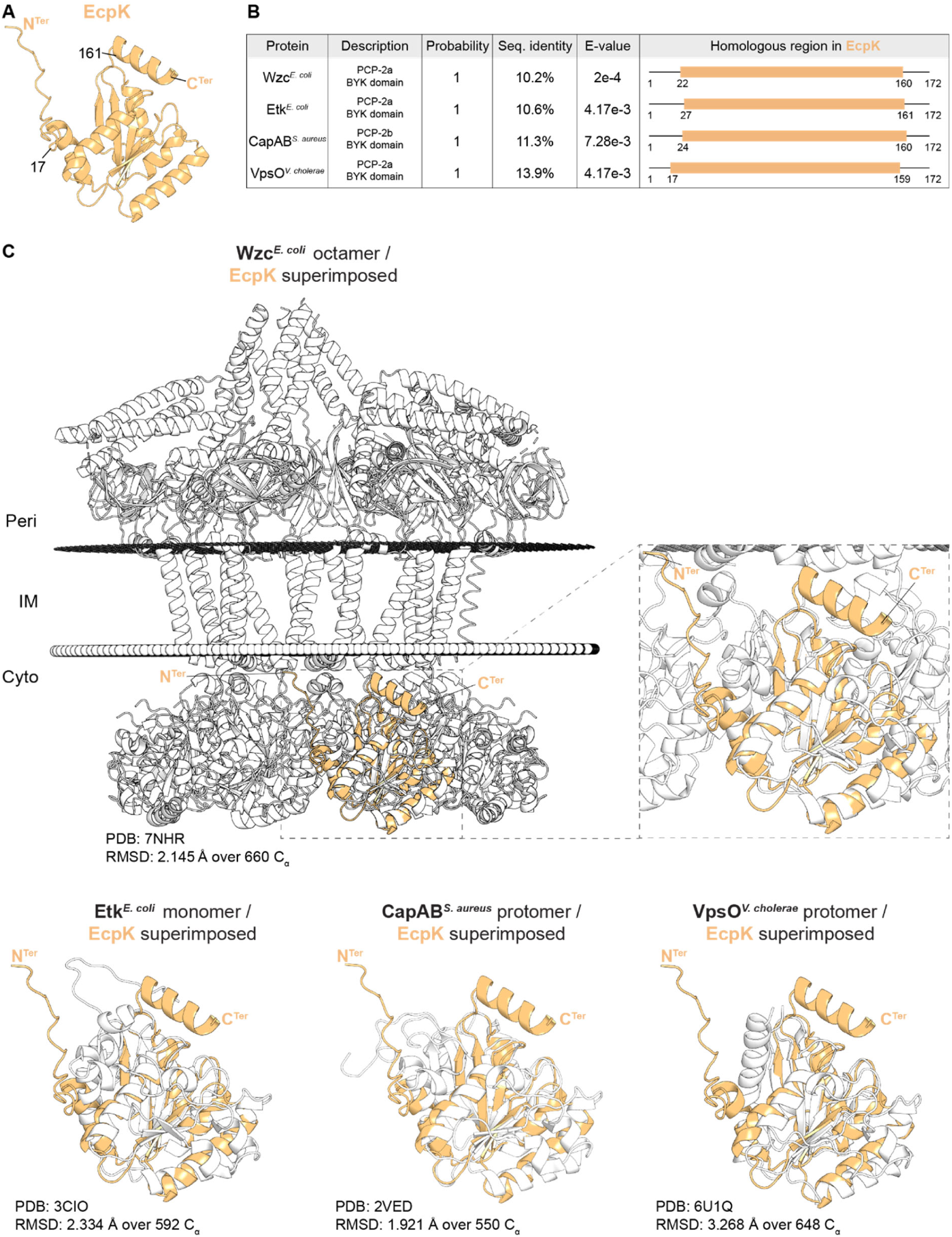
EcpK is structurally homologous to PCP-2 BYK domains. (A) AlphaFold2 structural model of EcpK, shown in light orange. The N- and C-termini are labeled, with residue numbers indicating the outermost boundaries of EcpK’s region homologous to the target structures in panel (B). (B) Foldseek analysis of EcpK against solved structures in PDB100. From left to right: Protein name, organism; description; probability; sequence identity; E-value; region of EcpK homologous to the target structure. Note that the CapAB*^S. aureus^* protein is CapB*^S. aureus^* fused to the C-terminal extension of CapA*^S. aureus^* (51). (C) Structural alignment of EcpK with Foldseek hits. EcpK is shown in light orange, with N- and C-termini marked. The target structures are shown in grey/white. Upper, left panel, alignment of EcpK with Wzc*^E. coli^* (32), with a root mean square deviation (RMSD) of 2.145 Å over 660 C_α_. The predicted position of the Wzc*^E. coli^* octamer within the IM was calculated using the PPM server (Lomize et al., 2022). Upper, right panel, zoomed in image of the Wzc*^E. coli^*/EcpK alignment. Lower panel, left, alignment of EcpK with Etk*^E. coli^* (50), with an RMSD of 2.334 over 592 C_α_. Lower panel, middle, alignment of EcpK with CapB*^S. aureus^* fused to the C-terminal extension of CapA*^S. aureus^* (51), with an RMSD of 1.921 over 550 C_α_. Lower panel, right, alignment of EcpK with VpsO*^V. cholerae^* (48), with an RMSD of 3.268 over 648 C_α_.

The active site of BYKs comprises variants of the Walker A and Walker B motifs, essential for nucleotide binding and hydrolysis, respectively (41, 43). Further, the Walker A’ motif facilitates recognition of Tyr-residues, interacts with the phosphate moiety of the bound nucleotide, and initiates phosphorylation (41, 43). These three motifs, as well as the Tyr-rich C-terminal tail of the BYK, are well conserved in Wzc*^E. coli^*, Etk*^E. coli^*, VpsO*^V. cholerae^* and CapB*^S. aureus^* (Fig. 2A). Strikingly, none of these motifs are present in EcpK (Fig. 2A) and no Tyr-residues are present in the C-terminal region of EcpK (Fig. 2A). Consistently, while the AlphaFold2 structural model of EcpK aligns with the active site of the Wzc*^E. coli^* BYK domain, the EcpK residues in this region do not match the residues of the Walker A, Walker B or Walker A’ motifs (Fig. 2B). Furthermore, whereas the C-terminus of Wzc*^E. coli^* is a flexible, Tyr-rich tail, the C-terminus of EcpK is an α-helix without Tyr-residues but instead comprises several Lys- and Arg-residues (Fig. 2C). Notably, all myxobacterial orthologs of EcpK lack the consensus Walker motifs and the Tyr-rich C-terminus (Fig. S3). Altogether, because EcpK is structurally similar to BYK domains, but lacks signature motifs necessary for catalytic BYK activity, we conclude that EcpK, along with its myxobacterial orthologs, is a BY pseudokinase.

**Figure 2.**
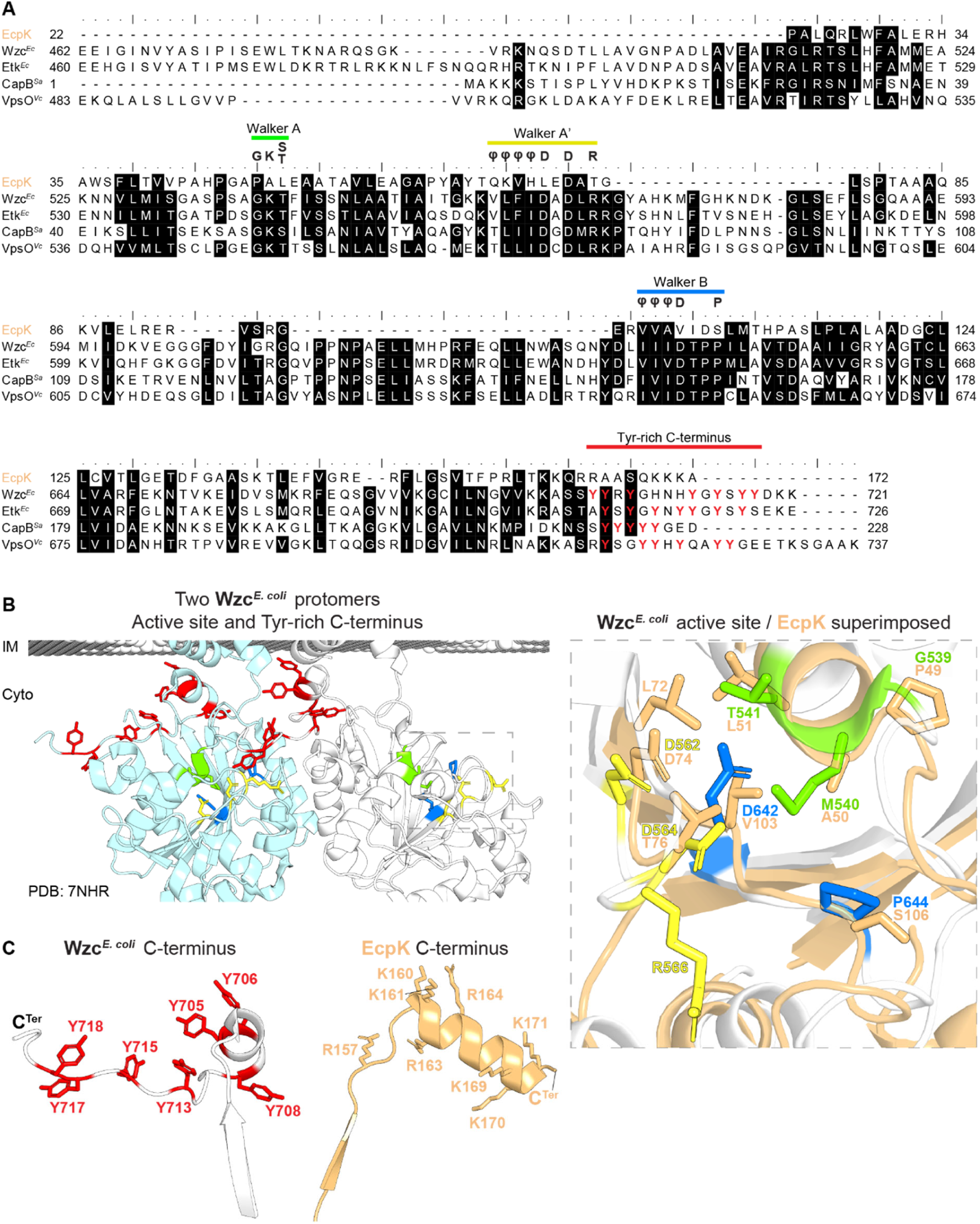
EcpK is a BY pseudokinase. (A) Sequence alignment of EcpK with the proteins identified in the Foldseek analysis in Fig. 1B and C. The BYK signature motifs (Walker A, Walker A’, Walker B) and the Tyr-rich C-terminal tail are indicated. Consensus Walker motif sequences (43) are shown, with φ representing hydrophobic residues. Amino acid residue numbering corresponds to the full-length sequences of the proteins. (B) Structural comparison of EcpK with the active site of Wzc*^E. coli^*. Left panel, base of the TMH and BYK domains of two Wzc*^E. coli^* protomers, shown in grey/white and light-blue. Walker motifs and Tyr-rich are colored as in (A). Right panel, active site of Wzc*^E. coli^* superimposed with EcpK. Amino acid residues corresponding to the Walker motifs in Wzc*^E. coli^* are shown and colored as in (A), with aligned residues in EcpK shown in light-orange. Note the amino acid substitution of the Walker A residue Lys^540^ to Met in Wzc*^E. coli^* in the non-phosphorylated, octameric structure (32). (C) Comparison of C-termini of Wzc*^E. coli^* and EcpK. Left panel, C-terminus of Wzc*^E. coli^*, with Tyr residues shown in red. Right panel, C-terminus of EcpK, with Lys and Arg residues shown in light orange.

### EcpK is important for EPS biosynthesis

To investigate the role of EcpK in EPS biosynthesis, we generated an in-frame deletion in *ecpK* (Δ*ecpK*). Using plate-based colorimetric assays with Trypan blue and Congo red as indicators of EPS production, we observed that the wild type (WT) strain DK1622 synthesized EPS (Fig. 3). By contrast, the Δ*ecpK* mutation led to a strong defect in EPS biosynthesis (Fig. 3), similarly to the Δ*epsZ* and Δ*epsV* mutations (Fig. 3) that disrupt genes encoding key proteins in the EPS biosynthetic pathway (Fig. S1D) (21, 64-66). Because EPS is important for T4P-dependent motility (63), we examined motility on 0.5% agar, which is favorable to T4P-dependent motility (69). The WT formed long flares at the colony edge characteristic of T4P-dependent motility (Fig. 3), while the Δ*pilA* mutant, which lacks the major subunit of T4P (70) and served as a negative control, generated a smooth-edged colony with limited expansion (Fig. 3). As previously shown (21, 64), the Δ*epsZ* and the Δ*epsV* mutants formed colonies with short flares and significantly reduced expansion compared to the WT, indicative of a defect in T4P-dependent motility (Fig. 3). Importantly, the Δ*ecpK* mutant also generated colonies with short flares and with significantly reduced colony expansion (Fig. 3). These findings are in agreement with previous reports, in which the Δ*ecpK* mutant was described as having a defect in EPS biosynthesis and T4P-dependent motility (66). On 1.5% agar, which is favorable to gliding motility (69), the WT displayed the single cells at the colony edge characteristic of gliding motility (Fig. 3), while the Δ*aglQ* mutant, which lacks a component of the Agl/Glt machinery for gliding (71, 72) and served as a negative control, did not. The Δ*ecpK,* Δ*pilA*, Δ*epsZ* and Δ*epsV* mutants all had single cells at the colony edges (Fig. 3). To rule out potential polar effects of the Δ*ecpK* mutation (Fig. S1A), we performed complementation experiments by ectopically expressing *ecpK* from its native promoter on a plasmid integrated in a single copy at the Mx8 *attB* site in the Δ*ecpK* mutant. Ectopic expression of *ecpK* fully restored EPS biosynthesis and T4P-dependent motility in the Δ*ecpK* mutant (Fig. 3).

**Figure 3.**
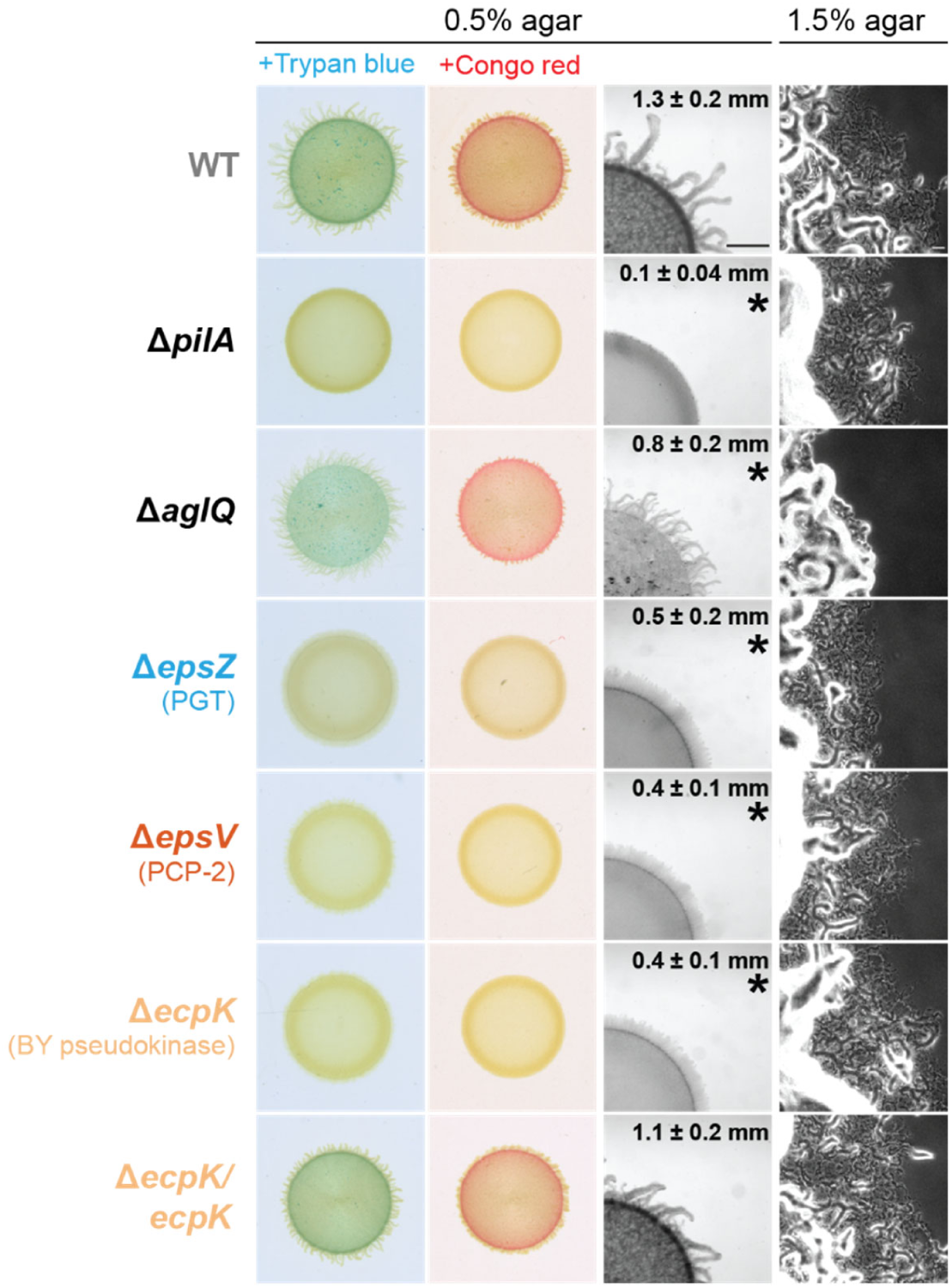
EcpK is important for EPS biosynthesis. Two left columns, EPS biosynthesis was assessed by spotting cells on 0.5% agar supplemented with 0.5% CTT and either Congo red or Trypan blue, with image recording after 24 h. Two right columns, T4P-dependent motility and gliding motility were analyzed on 0.5% and 1.5% agar, respectively, supplemented with 0.5% CTT, and images were recorded after 24 h. The Δ*epsZ* and the Δ*epsV* mutants, which lack components of the EPS biosynthetic pathway (64), were used as negative controls for EPS biosynthesis. For T4P-dependent motility assays, numbers indicate the colony expansion from the edge of the colony over 24 h as mean ± standard deviation (*n* = 3 biological replicates). *, *p* < 0.05, Welch’s test against WT. Scale bars, 1 mm (middle right) and 50 μm (right). For generating the in-frame deletion in *ecpK*, we reannotated *ecpK*. The previously predicted start codon (TTG) is unlikely in the GC-rich genome of *M. xanthus*. Instead, we identified an ATG codon 14 codons downstream as the likely start codon, supported by comparison of the N-terminal sequences of EcpK and its myxobacterial orthologs (Fig. S3).

We conclude that EcpK is critical for EPS biosynthesis and likely functions as an integral component of the Wzx/Wzy-dependent EPS biosynthetic pathway.

### EcpK and EpsV directly interact

Because EpsV lacks a BYK domain and EcpK is a BY pseudokinase, we speculated that EcpK could interact with EpsV to form a non-canonical bipartite PCP, and/or could interact with another component of the EPS biosynthetic pathway (Fig. S1D). Previous studies have shown that proteins that directly interact can mutually stabilize each other in *M. xanthus* (21, 73-76). Therefore, to identify potential EcpK interaction partner(s), we determined the accumulation level of *M. xanthus* proteins in cells grown on a solid surface using whole-cell label-free quantitative (LFQ) mass spectrometry-based proteomics.

In the whole-cell proteomics analysis, we initially focused on the Eps proteins (Fig. S1D). We detected all Eps proteins in the WT, except for the integral IM protein Wzy_EPS_ (Fig. 4A). Interestingly, in the Δ*ecpK* mutant, EpsV did not detectably accumulate, while the remaining Eps proteins had levels similar to those in the WT (Fig. 4A). Conversely, in the Δ*epsV* mutant, EcpK accumulated at significantly reduced levels (Fig. 4A). Additionally, accumulation of the GT EpsU was strongly reduced, and accumulation of the periplasmic ^D1D2^OPX protein EpsY, which is part of the bipartite EpsY/EpsX translocon (Fig. S1D), was slightly reduced (Fig. 4A). The remaining Eps proteins accumulated at levels comparable to those in the WT (Fig. 4A). Importantly, previous whole-cell proteomics analyses performed under the same conditions as those used here, showed that EcpK and EpsV accumulated at WT levels in the Δ*epsZ* mutant (21), strongly suggesting that lack of EPS biosynthesis *per se* does not cause the reduced accumulation of EpsV in the Δ*ecpK* mutant and *vice versa*. These findings suggest that the accumulation of EcpK and EpsV mutually depends on each other.

**Figure 4.**
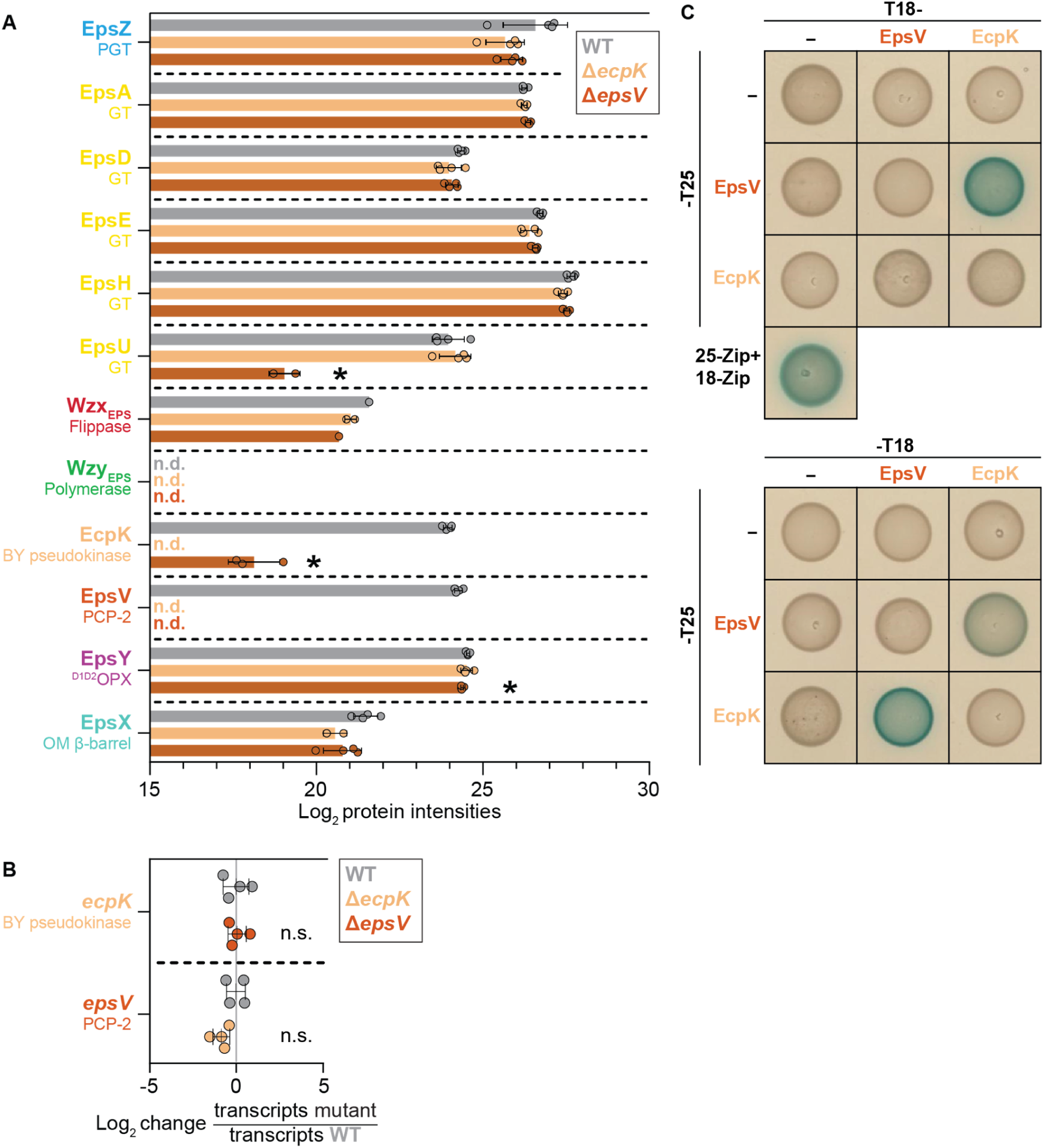
EcpK interacts directly with EpsV. (A) EcpK and EpsV mutually stabilize each other. Protein amounts in whole-cell proteomes of *M. xanthus* strains were quantified using LFQ mass spectrometry-based proteomics. Normalized Log_2_ intensities of Eps proteins in the indicated strains are shown. Each data point represents a biological replicate (*n* = 4 biological replicates). Error bars indicate standard deviation across these replicates. *, *p* < 0.01, Welch’s test against WT. "n.d." indicates that a protein was not detected in any replicate in a strain, and is shown in the corresponding strain’s color. (B) RT-qPCR analysis of *ecpK* and *epsV* transcripts levels. Transcript levels of *ecpK* and *epsV* were assessed using RT-qPCR and normalized to the expression of the internal reference gene *MXAN_3298*, encoding elongation factor Tu (73). Data are presented as Log_2_-fold change in transcript levels in the mutant strains relative to the WT. Data points represent biological replicates (*n* = 4 biological replicates, each with two technical replicates) and are colored according to the strain analyzed. The center marker and error bars represent mean and standard deviation, respectively. Statistical significance was evaluated using Welch’s test against WT, with "n.s." indicating no significant difference (*p* < 0.01). (C) EcpK and EpsV interact in BACTH assay. Full-length EcpK and EpsV were fused to the N- and C-termini of T18 and to the N-terminus of T25, respectively. The positive control T25-Zip + T18-Zip is shown in the lower left corner. T18 and T25 alone served as negative controls. Shown are representative images of *E. coli* BTH101 expressing the indicated protein fusions. For each tested interaction pair, at least two clones were analyzed with similar results.

Next, we considered that EcpK could also be involved in regulating the accumulation of regulators of EPS biosynthesis (63). However, except for TmoK, which by an unknown mechanism acts as an inhibitor of EPS biosynthesis and was slightly but significantly reduced, we found no significant changes in the levels of these regulators in the Δ*ecpK* mutant compared to the WT (Fig. S4).

To investigate whether the observed changes in EpsV accumulation in the Δ*ecpK* mutant and *vice versa* were caused by transcriptional changes, we performed quantitative reverse-transcriptase PCR (qRT-PCR). As in the whole-cell LFQ mass spectrometry-based proteomics experiments, cells were grown on a solid surface. In the Δ*ecpK* mutant, *epsV* was transcribed at WT levels (Fig. 4B); similarly, in the Δ*epsV* mutant, *ecpK* was transcribed at WT levels (Fig. 4B). These findings demonstrate that the mutual accumulation dependency of EcpK and EpsV (Fig. 4A) occurs at a post-transcriptional level, supporting that EcpK and EpsV interact, thereby stabilizing each other.

Finally, to investigate whether EcpK and EpsV interact directly, we performed a bacterial adenylate cyclase-based two-hybrid (BACTH) assay. In this experiment, we observed a direct interaction between EcpK and EpsV (Fig. 4C), while we observed neither EcpK nor EpsV self-interaction (Fig. 4C). Altogether, these results strongly support that EcpK specifically activates EPS biosynthesis through direct interaction with EpsV.

### EcpK may directly interact with the cytoplasmic base of EpsV

To investigate how EcpK and EpsV could interact directly, we generated a model of a EcpK/EpsV heterocomplex using AlphaFold-Multimer (77). As in the AlphaFold2 model of monomeric EpsV (21), the resulting high-confidence model (Fig. S5) supports that EpsV has two TMHs, a periplasmic domain rich in α-helices, and short N- and C-terminal cytoplasmic extensions of 37 and 25 residues, respectively (Fig. 5A). In the heterocomplex, EcpK is placed at the cytoplasmic base of EpsV (Fig. 5A and B). EcpK and EpsV interact *via* the C-terminal cytoplasmic extension of EpsV, which folds into a groove on EcpK’s membrane-facing side (Fig. 5C). This groove ends in two negatively charged pockets in EcpK in which two Arg residues in EpsV’s C-terminus anchor in a hook-like arrangement (Fig. 5C). Additionally, most of EcpK’s surface that faces the IM’s cytoplasmic leaflet is positively charged, suggesting electrostatic interactions with the negatively charged IM (Fig. 5C). This positive charge is partly due to EcpK’s C-terminal α-helix, which contains numerous Lys- and Arg-residues (Fig. 5A−C, Fig. 2A and 2C), a characteristic conserved among several myxobacterial EcpK orthologs (Fig. S3). Altogether, this structural model further supports a direct interaction between EcpK and EpsV.

**Figure 5.**
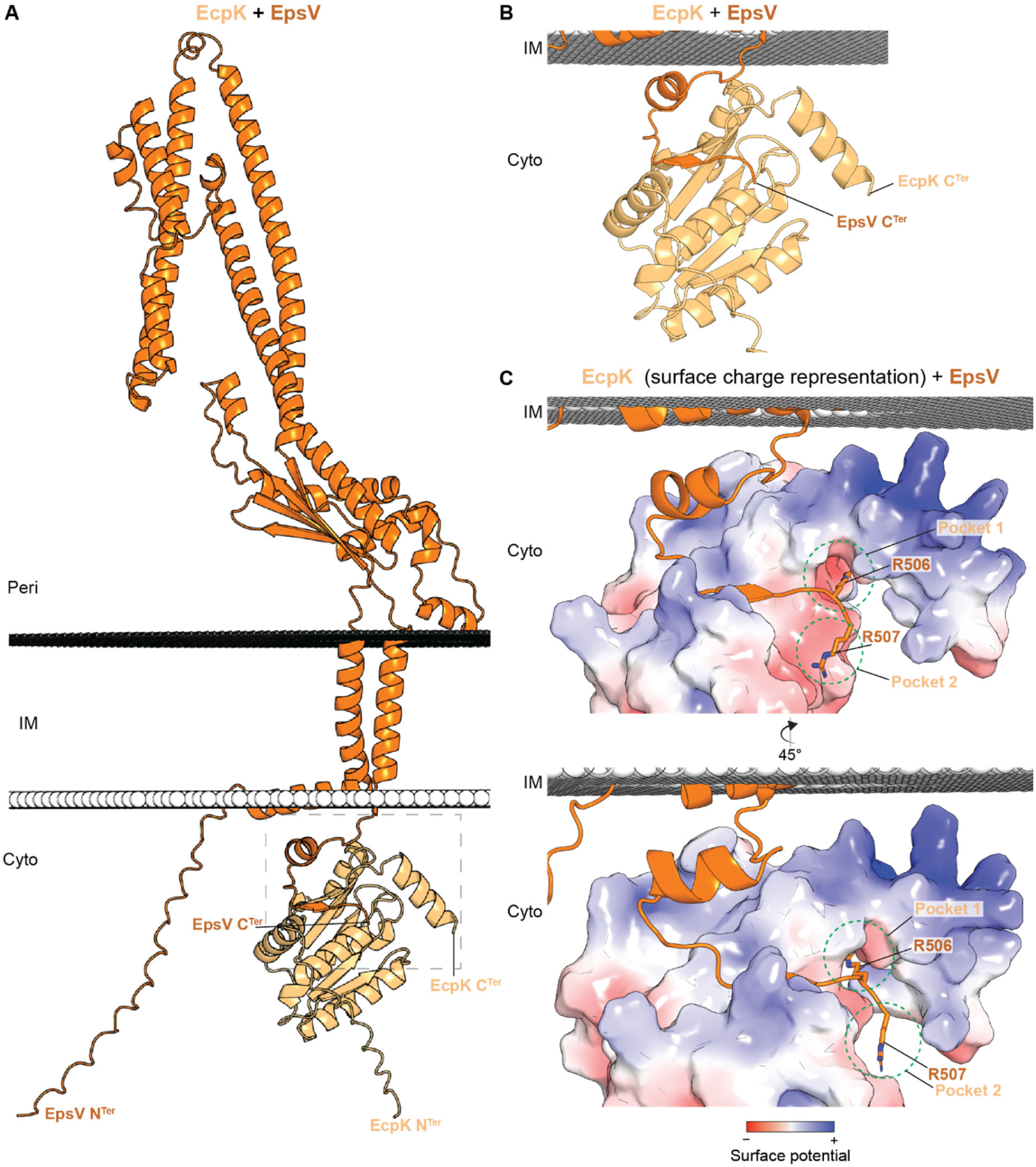
Computational structural characterization of the EcpK-EpsV complex. (A) AlphaFold2-Multimer model of the EcpK/EpsV complex. EcpK is shown in light orange, and EpsV in orange. The predicted position of the EcpK-EpsV complex within the membrane was calculated using the PPM server (107). Model rank 1 is shown. (B) Zoomed in image of the EcpK/EpsV interaction involving the C-terminal cytoplasmic extension of EpsV that folds into a groove on EcpK’s membrane-facing side. (C) Zoomed in image of the EcpK/EpsV interaction with EcpK shown in surface charge representation (contoured from +5 to −5 kT e^−1^), which was computed using pdb2pqr *via* the Adaptive Poisson-Boltzmann Solver server (106). Negative and positive charges are colored red and blue, respectively. The two Arg residues at EpsV’s C-terminus are highlighted together with the two negatively charged pockets in EcpK in which the Arg residues anchor in a hook-like arrangement. Note that most of EcpK’s surface facing the IM’s cytoplasmic leaflet is positively charged.

The BYK domain of Wzc*^E. coli^*, both when expressed on its own and in the context of the full-length protein, and the CapB*^S. aureus^* BYK fused to the C-terminal extension of CapA*^S. aureus^* form octamers in the non-phosphorylated state (32, 49, 51). To explore whether the EcpK/EpsV complex could similarly adopt an octameric structure, we used AlphaFold2-Multimer (77) to model eight copies of EcpK together with eight copies of EpsV’s cytoplasmic C-terminal 25 residues (residues 483-507). Interestingly, the resulting model predicted EcpK/EpsV^483-507^ in an octameric ring with partially good confidence (Fig. S6). As in the heterocomplex comprising each one copy of EcpK and full-length EpsV (Fig. 5A-C), the EpsV cytoplasmic C-terminal 25 residues fold into the EcpK groove in all eight pairs of protomers, while the positively charged C-terminal α-helices in the eight EcpK protomers are disconnected from the ring structure and do not engage in protein-protein interactions (Fig. S6). This model further support that EcpK and EpsV may interact to form an octameric complex.

## Discussion

In this study, we identify EcpK as a structural homolog of BYK domains of PCP-2 proteins. However, EcpK lacks conserved Walker motifs for nucleotide binding, hydrolysis and Tyr phosphorylation, as well as the Tyr-rich C-terminal tail. Based on these findings, we conclude that EcpK is a BY pseudokinase. Our experimental evidence document that EcpK directly interacts with the PCP-2 protein of the EPS biosynthetic pathway, EpsV, which lacks a BYK domain. By means of computational modeling, we located the EcpK/EpsV interaction interface to the cytoplasmic C-terminus of EpsV, which in our structural model folds into a groove on EcpK’s IM-facing surface. Based on these lines of evidence and because EcpK is essential for EPS biosynthesis, we conclude that the EcpK BY pseudokinase is a novel component of the EPS biosynthetic pathway and stimulates EPS biosynthesis by direct interaction with EpsV, together forming a non-canonical bipartite PCP-2.

Based on sequence analysis, we predict that the EpsV and EcpK orthologs in other myxobacteria interact and function similarly to the EcpK/EpsV pair. Moreover, a sequence-based analysis suggests that similar protein pairs are present in Wzx/Wzy-dependent pathways of several other Gram-negative bacteria (25). One of these systems has been experimentally analyzed, i.e. HfsAB of the Wzx/Wzy-dependent pathway for holdfast biosynthesis in *C. crescentus*. Here, the stand-alone BY pseudokinase HfsB lacks catalytic residues for Tyr kinase activity and the C-terminal Tyr-rich tail, the PCP-2 HfsA lacks a BYK domain, and both are essential for holdfast biosynthesis (62, 78, 79). Interestingly, HfsB is important for the stability of HfsA (62), suggesting that HfsB and HfsA, similar to EcpK and EpsV, form a non-canonical bipartite PCP-2. Indeed a high confidence AlphaFold-Multimer model of the HfsB/HfsA heterocomplex suggests that the last 27 C-terminal, cytoplasmic residues of HfsA interact with HfsB (Fig. S7 and 8A-C). In the models of the EcpK/EpsV and HfsB/HfsA heterocomplexes, EcpK and HfsB differ in the predicted interactions of their C-terminal α-helices. Specifically, EcpK’s C-terminal α-helix is positively charged and may associate with the IM, whereas HfsB’s C-terminal α-helix is predicted to interact with the base of one of HfsA’s two TMHs. The EcpK/EpsV and HfsB/HfsA interactions are overall similar to the interaction between CapA and CapB in *S. aureus* that involves the 29 residues cytoplasmic C-terminal extension of CapA (34, 51). Altogether, these findings support the idea that in both pathways, the BY pseudokinase functions through direct interaction with its PCP-2 partner protein.

It is well established that PCP-2 proteins comprise two subfamilies, i.e. Wzc*^E. coli^*-like PCP-2a proteins, in which the transmembrane/periplasmic part and the BYK domain are joined in a single polypeptide, and the CapAB*^S. aureus^*-like PCP-2b proteins, in which the transmembrane/extracytoplasmic part and the BYK domain are separate proteins. Based on the characterization of the EcpK/EpsV and HfsB/HfsA pairs, we suggest the existence of a third subfamily, comprising bipartite PCP-2 proteins, in which the transmembrane/periplasmic part partners with a stand-alone BY pseudokinase important for function. Based on the analysis of Cuthbertson *et al.* (25), this subfamily is likely present in Wzx/Wzy-dependent pathways of many other Gram-negative bacteria. We note that based on the Cuthbertson *et al.* analysis, similar PCP-2 variants occur, in which the transmembrane/periplasmic part and the BY pseudokinase are joined in a single polypeptide.

In the current model, PCP-2 proteins functionally depend on cycles of octamer assembly/disassembly, with these cycles being controlled by the phosphorylation level of the Tyr-rich tail of the associated BYK domains (32, 48, 49, 51). This brings up the question how a BY pseudokinase can stimulate the function of a PCP-2. In eukaryotes, Tyr pseudokinases function by forming heterodimers with active kinases, e.g. the catalytically inactive HER3 pseudokinase heterodimerizes with the catalytically active receptor Tyr kinases HER1 and HER2 to stimulate kinase activity (80, 81). Alternatively, they can negatively regulate Tyr kinase activity by an intramolecular mechanism, e.g. in Janus kinases, the Tyr pseudokinase JH2 domain inhibits the Tyr kinase activity of the catalytically active JH1 domain (82). *M. xanthus* encodes two *bona fide* BYKs (83, 84). In the Wzx/Wzy-dependent pathway for the spore coat polysaccharide (SPS), the catalytically active stand-alone BYK BtkA (also referred to as ExoD (85)), which is essential for SPS biosynthesis, features Walker motifs, a Tyr-rich C-terminus, and undergoes Tyr phosphorylation in the presence of the PCP-2 partner ExoC (83). A Δ*btkA* mutant synthesizes WT levels of EPS (83). The Wzx/Wzy-dependent pathway for the biosurfactant polysaccharide (BPS) includes the Wzc*^E. coli^*-like PCP-2a protein BtkB (also referred to as WzcB (66)), in which the BYK domain is catalytically active and contains the conserved Walker motifs, a Tyr-rich C-terminus, and undergoes Tyr phosphorylation (84). The Δ*btkB* mutant phenocopies mutants lacking other components of the BPS biosynthetic pathway (66) and has been reporter to have either a slight defect in EPS biosynthesis (64, 84), which was suggested to be caused by sequestration of Und-P, or WT-levels of EPS biosynthesis (66). Also, lack of the phospho-tyrosine phosphatase PhpA, which dephosphorylates Tyr-phosphorylated BtkA and BtkB (86), causes a slight increase in EPS biosynthesis (86). These observations suggest that neither BtkA, BtkB nor PhpA directly contribute to EPS biosynthesis, supporting that EcpK, and by implication other BY pseudokinases involved in polysaccharide biosynthesis, do not functionally depend on oligomerization or interaction(s) with catalytically active BYKs.

Based on these considerations, we suggest that BY pseudokinases such as EcpK and HfsB function as scaffolds for the transmembrane/periplasmic part of the PCP-2 partner protein to facilitate PCP-2 function independently of tyrosine-phosphorylation. In this model, we speculate that EpsV and HfsA would depend on octamer assembly/disassembly cycles for function, and that EcpK and HfsB would guide these cycles by binding/unbinding cycles independently of BYK-dependent phosphorylation and phosphatase-dependent dephosphorylation cycles.

Altogether, this work provides a framework for future biochemical and structural studies of EcpK and EpsV as well as of other Wzx/Wzy-dependent pathways for polysaccharide biosynthesis containing a BY pseudokinase.

## Supporting information

All Supplementary Information

## Acknowledgement

The authors thank Dobromir Szadkowski for providing a Matlab script to help with illustrating the AlphaFold model confidence plots and Anke Treuner-Lange and Marco Herfurth for help in AlphaFold-Multimer modeling. The Max Planck Society supported this work.

## Data Availability

The data supporting the findings of this study are all included in the manuscript and its supplementary file. The mass spectrometry proteomics data of whole cell proteomics have been deposited to the ProteomeXchange Consortium via the PRIDE partner repository with the dataset identifier PXD058227. All materials are available from the corresponding author upon request.

## Materials and Methods

### Strains and cell growth

All *M. xanthus* strains used in this study are derivatives of the WT strain DK1622 (87) and are listed in Table 1. Plasmids and oligonucleotides are listed in Table 2 and Table S1, respectively. In-frame deletions were constructed by two-step homologous recombination following the protocol in (88). The complementation plasmid was integrated in a single copy by site-specific recombination at the Mx8 *attB* site. All plasmids were verified by DNA sequencing, and all strains were verified by PCR. *M. xanthus* cultures were grown at 32°C in 1% CTT broth (1% [wt/vol] Bacto casitone, 10 mM Tris-HCl [pH 8.0], 1 mM K_2_HPO_4_/KH_2_PO_4_ [pH 7.6], 8 mM MgSO_4_) or on 1.5% agar supplemented with 1% CTT and kanamycin (50 μg mL^−1^) or oxytetracycline (10 μg mL^−1^) when appropriate (89). Plasmids were propagated in *E. coli* NEB Turbo at 37°C in lysogeny broth (LB) (90) supplemented with kanamycin (50 μg mL^−1^), tetracycline (20 μg mL^−1^) or carbenicillin (100 μg mL^-1^) when required.

**Table 1.** Strains used in this work.

| Species and strain | Genotype | Reference or source |
| --- | --- | --- |
| *M. xanthus* |  |  |
| DK1622 | WT | (87) |
| DK10410 | $\Delta pilA$ | (108) |
| SA5923 | $\Delta aglQ$ | (76) |
| SA7400 | $\Delta epsZ$ | (64) |
| SA7406 | $\Delta epsV$ | (64) |
| SA11618 | $\Delta ecpK$ | This study |
| SA11623 | $\Delta ecpK attB::pLBL004$ | This study |
| *E. coli* |  |  |
| *E. coli* NEB Turbo | F' *proA*<sup>+</sup>*B*<sup>+</sup> *lacI*<sup>q</sup> $\Delta lacZM15$ / *fhuA2* $\Delta(lac-proAB)$ *glnV galK16 galE15 R(zgb-210::Tn10)*Tet<sup>S</sup> *endA1 thi-1* $\Delta(hsdS-mcrB)5$ | New England Biolabs |
| BTH101 | F<sup>-</sup> *cya-99 araD139 galE15 galK16 rpsL1* (Str<sup>R</sup>) *hsdR2 mcrA1 mcrB1* | Euromedex |

**Table 2.**
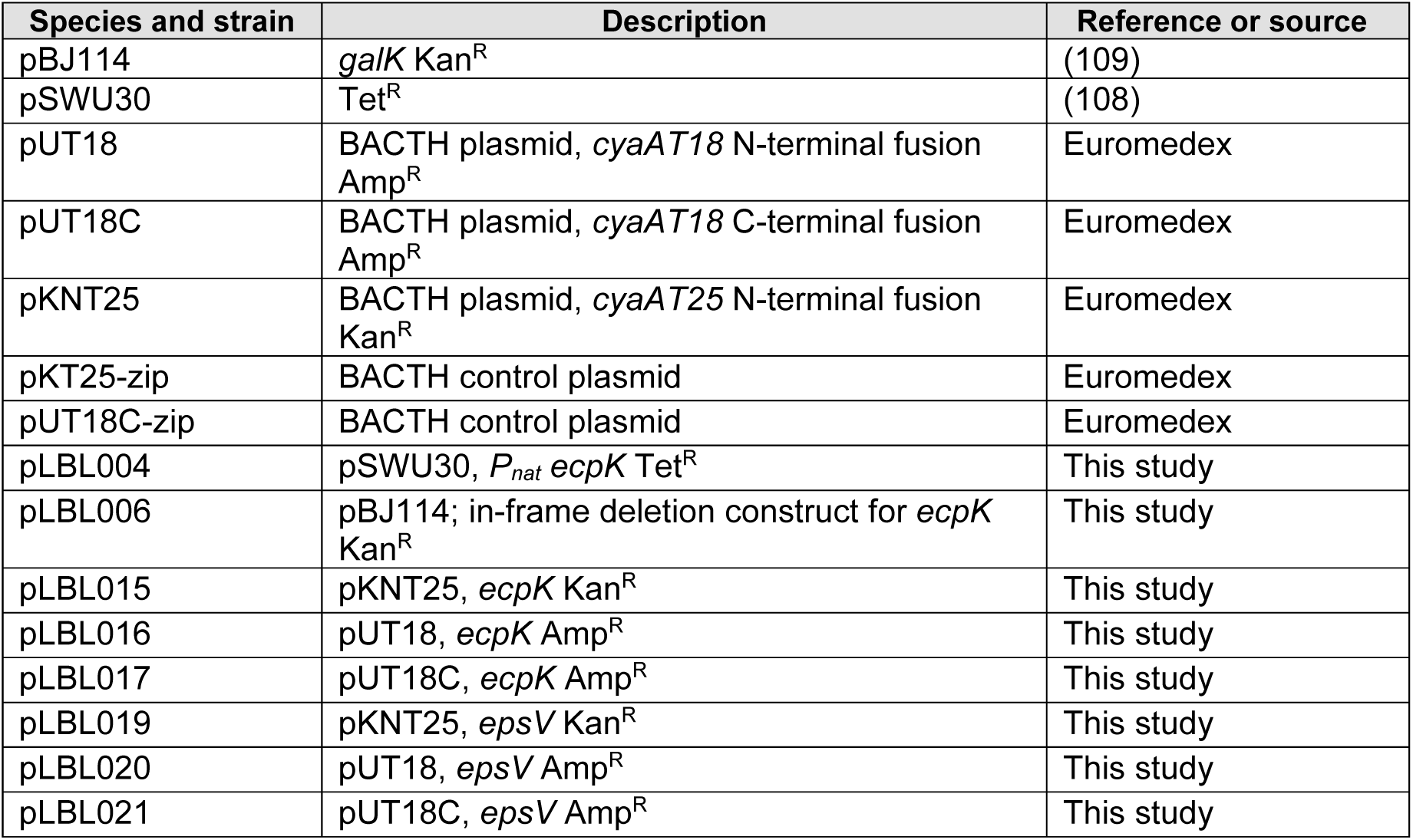
Plasmids used in this work.

### Plasmid construction

For pLBL006 (for generating the in-frame deletion in *ecpK*), AB and CD fragments were amplified from genomic DNA of DK1622 using the primer pairs LB1/LB40 and LB18/LB4, respectively. Subsequently, the AB and CD fragments were used as templates for an overlap PCR with the primer pair LB1/LB4 to generate the AD fragment. The AD fragment was cloned into pBJ114 *via* KpnI/XbaI sites. For pLBL004 (for generating a strain ectopically expressing from the *attB* site *ecpK* from the native promoter), P_nat_ *ecpK* was amplified from genomic DNA of DK1622 using the primer pair LB9/LB10. The fragment, which comprises *ecpK* and 542 bp upstream of its start codon, was cloned into pSWU30 *via* HindIII/XbaI sites.

For constructing BACTH plasmids, all DNA fragments were amplified from *M. xanthus* genomic DNA. For pLBL015 and pLBL016, full-length *ecpK* was amplified using the primer pair LB59/LB62 and cloned into pKNT25 and pUT18, respectively, *via* HindIII/KpnI sites. For pLBL017, full-length *ecpK* was amplified using the primer pair LB60/LB62 and cloned into pUT18C *via* XbaI/KpnI sites. For pLBL019 and pLB020, full-length *epsV* was amplified using the primer pair LB61/LB65 and cloned into pKNT25 and pUT18, respectively, *via* HindIII/XbaI sites. For pLBL021, full-length *epsV* was amplified using primer the primer pair LB63/LB64 and cloned into pUT18C *via* XbaI/KpnI sites.

### Detection of EPS biosynthesis

Colony-based colorimetric EPS assays were conducted as described in (64). Exponentially growing cells in suspension cultures were sedimented by centrifugation (3 min, 6,000 × *g* at room temperature) and resuspended in 1% CTT to a final cell density of 7 × 10^9^ cells mL^−1^. Next, 20 μL of this suspension was spotted onto 0.5% agar plates containing 0.5% CTT supplemented with either 10 μg mL^−1^ Trypan blue or 20 μg mL^−1^ Congo red. Plates were incubated at 32°C and imaged after 24 h.

### Motility assays

Motility assays were performed following the method described in (69). Exponentially growing cells in suspension cultures were sedimented by centrifugation (3 min, 6,000 × g, room temperature) and resuspended in 1% CTT medium to a final cell density of 7 × 10⁹ cells mL^−1^. 5 µL aliquots of this cell suspension were placed onto 0.5% and 1.5% agar supplemented with 0.5% CTT, followed by incubation at 32°C for 24 h. Imaging was performed using a M205FA stereomicroscope (Leica Microsystems, Wetzlar, Germany) equipped with a Hamamatsu ORCA-Flash V2 digital CMOS camera (Hamamatsu Photonics, Hersching, Germany) and a DMi8 inverted microscope with a DFC9000 GT camera (both Leica Microsystems, Wetzlar, Germany).

### Proteomic analysis using data independent acquisition-mass spectrometry (DIA-MS)

Whole-cell proteomics experiments of cells grown on a solid surface were performed as described (21). Briefly, cells were grown on 1.5% agar plates supplemented with 1% CTT for 72 h at 32°C. Next, 35 mg per sample were harvested and washed twice in 0.5 mL 1× phosphate-buffered saline (PBS) (137 mM NaCl, 2.7 mM KCl, 10 mM Na_2_HPO_4_, 1.8 mM KH_2_PO_4_, pH 7.5) supplemented with 2× protease inhibitor (Roche). The cells were sedimented and resuspended in 0.2 mL 0.1 M ammonium bicarbonate containing 2% (w/vol) sodium lauroyl sarcosinate (SLS), followed by incubation at 95°C for 1 h. Next, the samples were centrifuged at 14,000× *g* for 5 min, and the supernatant harvested. Next, 1.2 mL freezer-cold acetone was added to the supernatant, mixed, and incubated at −80°C for at least 2 h. Next, the samples were centrifuged at 21,000× *g* for 15 min at 4°C. The supernatant was discarded, and the pellet was washed thrice with freezer-cold methanol. Next, the pellet was dried, and the methanol completely removed. The protein pellet was resuspended in 200 μL 0.5% SLS (w/vol), and the protein amount determined by bicinchoninic acid-based protein assay (Thermo Fisher Scientific). Proteins were reduced with 5 mM Tris(2-carboxyethyl) phosphine (Thermo Fisher Scientific) at 90°C for 15 min and alkylated using 10 mM iodoacetamid (Sigma Aldrich) at 25°C for 30 min in the dark. 50 µg protein was digested by 1 µg trypsin (Serva) at 30°C overnight.

After digestion, SLS was precipitated by acidification and peptides were desalted by using C18 solid phase extraction cartridges (Macherey-Nagel). Cartridges were prepared for sample loading by adding acetonitrile (ACN), followed by 0.1% trifluoroacetic acid (TFA, Thermo Fisher Scientific). Peptides were loaded on equilibrated cartridges, washed with 5% ACN/0.1% TFA containing buffer and finally eluted with 50% ACN and 0.1% TFA.

Dried peptides were reconstituted in 0.1% TFA and then analyzed using LC-MS carried out on an Ultimate 3000 RSLC nano connected to an Exploris 480 Mass Spectrometer *via* a nanospray flex ion source (all Thermo FisherScientific) and an in-house packed HPLC C18 column (75 μm × 42 cm). The following separating gradient was used: 94% solvent A (0.15% formic acid) and 6% solvent B (99.85% ACN, 0.15% formic acid) to 25% solvent B over 95 minutes at a flow rate of 300 nL/min, followed by an additional increase of solvent B 35% over 25 min.

MS raw data was acquired in data independent acquisition mode. In short, spray voltage was set to 2.3 kV, funnel radio frequency level at 40 and the ion transfer capillary heated to 275°C. For DIA experiments full MS resolutions were set to 120,000 at m/z 200 and full MS, AGC (Automatic Gain Control) target was 300% with an 50 ms IT (Ion Accumulation Time). Mass range was set to 350–1400. AGC target value for fragment spectra was set at 3000%. 45 windows of 14 Da plus 1 Da overlap were used. Resolution was set to 15,000 and MS/MS IT to 22 ms. Stepped HCD (high energy collision dissociation) collision energy of 25, 27.5, 30 % was used. MS1 data was acquired in profile, MS2 DIA data in centroid mode.

For analyzing DIA data the neural network (NN) based DIA-NN suite version 1.8 (91) and an Uniprot protein database for *M. xanthus* was used. A data set centric spectral library for the DIA analysis was generated. DIA-NN performed noise interference correction (mass correction, RT prediction and precursor/fragment co-elution correlation) and peptide precursor signal extraction of the DIA-NN raw data. The following parameters were used: Full tryptic digest was allowed with two missed cleavage sites, and oxidized methionines and carbamidomethylated cysteines as modifications. Match between runs and remove likely interferences were enabled. The NN classifier was set to the single-pass mode, and protein inference was based on genes. Quantification strategy was set to any LC (high accuracy). Cross-run normalization was set to RT-dependent. Library generation was set to smart profiling. DIA-NN outputs were further evaluated using the SafeQuant (92, 93) script modified to process DIA-NN outputs.

The mass spectrometry proteomics data of whole cell proteomics experiments have been deposited to the ProteomeXchange Consortium (94) *via* the PRIDE (95) partner repository with the dataset identifier PXD058227.

### RT-qPCR

*M. xanthus* cells of the respective strains were grown in biological quadruplicates as described for the whole-cell proteomics experiment. Total RNA was extracted using the Monarch® Total RNA Miniprep Kit (New England Biolabs, Frankfurt, Germany). Briefly, 10^9^ cells were collected from the agar plates, resuspended in 200 μL lysis buffer (100 mM Tris-HCl [pH 7.6], 1 mg mL^−1^ lysozyme), and incubated at 25°C for 5 min. The manufacturer’s protocol was followed to purify RNA. Next, Turbo DNase (Thermo Scientific, Dreieich, Germany) was added to the RNA following the manufacturer’s protocol and subsequently removed using the Monarch® RNA Cleanup Kit (50 μg; New England Biolabs, Frankfurt, Germany). The LunaScript RT supermix kit (New England Biolabs, Frankfurt, Germany) was used to generate complementary DNA (cDNA) using 1 μg RNA. qPCRs were performed with two technical replicates per biological replicates on an Applied Biosystems 7500 real-time PCR system using the Luna universal qPCR master mix (New England Biolabs, Frankfurt, Germany) with the primers listed in Table S1. Differential gene expression analysis was performed following the comparative threshold cycle (*C_T_*) method (96). *MXAN_3298*, encoding the elongation factor Tu, was used as an internal reference gene, as described previously (73).

### Bacterial two-hybrid assay

BACTH experiments were conducted following the manufacturer’s (Euromedex) protocol. DNA fragments encoding *ecpK* and *epsV* were cloned into the vectors pUT18, pUT18C and pKNT25 to generate in-frame N- and C-terminal fusions with the T18 fragment of *Bordetella pertussis* adenylate cyclase CyaA, and N-terminal fusions with the T25 fragment of CyaA, respectively. Next, the indicated plasmid combinations (20 ng per plasmid) were co-transformed into competent *E. coli* BTH101 cells and incubated at 32°C for 24 h. As a positive control, pKT25-zip and pUT18C-zip plasmids, which encode leucine zipper motifs that dimerize, were co-transformed into competent *E. coli* BTH101 cells. As an indicator of protein-protein interaction, cyclic AMP (cAMP) synthesis by reconstituted CyaA was assessed qualitatively by observing blue color formation of colonies grown on LB agar plates containing appropriate antibiotics, 40 µg mL⁻¹ 5-bromo-4-chloro-3-indolyl-β-D-galactopyranoside (X-Gal), and 0.5 mM isopropyl-β-D-thiogalactopyranoside (IPTG). Three colonies harboring each of the indicated plasmid combinations were used to inoculate 200 µL LB with appropriate antibiotics and incubated at 32°C with shaking for 3 h. From each culture, 2 µL were spotted onto LB agar plates containing appropriate antibiotics, 40 µg mL^−1^ X-Gal, and 0.5 mM IPTG. These plates were incubated at 32°C for 24 hours, then transferred to 4°C and incubated for an additional 48 hours before imaging.

### Bioinformatics

Gene and protein sequences were obtained from the databases of KEGG (Kanehisa & Goto, 2000) or UniProt (97). The phylogenetic tree of Myxobacteria was generated following a workflow described in (64, 98) in MEGA-X (99) using the neighbor-joining method (100) and the genome sequences listed in Table S2. The reciprocal BLASTP hit method, following a workflow described in (Pérez-Burgos et al., 2020a, Pérez-Burgos et al., 2020b), was used for synteny analyses. Briefly, orthologs of an *M. xanthus* gene of interest in myxobacterial genomes listed in Table S2 were identified using the KEGG Sequence Similarity DataBase (101). Genes within a distance of <10 were considered in the same cluster and colored with identical colors. Clusters were considered separate when they had a distance of at least ten genes from another cluster. Genes not found in any cluster were considered orphan. Sequence alignments were computed using Clustal-Omega in MEGA-X (99).

Structure predictions were performed with AlphaFold2 and AlphaFold2-Multimer_v3 modeling *via* ColabFold (67, 77, 102) using the Alphafold2_mmseqs2 notebook with default settings. To evaluate AlphaFold-generated models, predicted local distance difference test (pLDDT) and predicted alignment error (pAE) graphs of five models were made using a custom-made Matlab R2020a (The MathWorks) script. These models were ranked based on combined pLDDT, pAE values and predicted (interface) template modeling score (p[i]TM) scores, with the best-ranked models used for further analysis and presentation. Per-residue model confidence was estimated based on pLDDT values (>90, high accuracy; 70 to 90, generally good accuracy; 50 to 70, low accuracy; <50, should not be interpreted) (67). The relative positioning of residues was validated by assessing pAE, measured in Å. The pAE graphs indicate the expected position error at residue X if the predicted and true structures were aligned on residue Y; the lower the pAE value, the higher the accuracy of the relative position of residue pairs and, consequently, the relative position of domains/subunits/proteins (67). piTM values (77) were further used to evaluate interface accuracy in multimeric models. A piTM score above 0.8 indicates an accurate interface, scores between 0.6 and 0.8 represent a grey zone where predictions may or may not be correct, and scores below 0.6 indicate a failed prediction (103, 104). The PAE Viewer was used to fetch average pLDDT values (105).

PyMOL (The PyMOL Molecular Graphics System, Version 2.4.1 Schrödinger, LLC) was used to analyze and visualize the structural models. For generating models colored based on pLDDT values, a custom command line was used (spectrum b, red_yellow_green_cyan_blue, minimum=50, maximum=90). Structural superimpositions were performed using the “super” method within the PyMOL Alignment plugin with default settings. For the calculation of surface charges, protein models were prepared using pdb2pqr with default settings (106), and electrostatics were calculated *via* the Adaptive Poisson-Boltzmann Solver (APBS) (106) plugin in PyMOL with default settings. Foldseek (68) was used to identify protein homologs in the PDB. For predicting the positioning of protein structural models within the IM, the PPM 3.0 web server (107) was used with default settings and type of membrane set to “Gram-negative bacteria inner membrane”.

### Statistical analysis

Statistics were performed using Welch’s tests. Data shown for T4P-dependent motility, gliding motility and EPS assays were obtained in at least three independent experiments with similar results. For LFQ-MS proteome analyses, four biological replicates were analyzed. For RT-qPCR, four biological replicates with each two technical replicates were analyzed. For qualitative determination of protein-protein interactions using the BACTH assay, similar results were obtained with at least two clones per each combination.

