## Supplementary material for "Identification of EcpK, a bacterial tyrosine pseudokinase important for exopolysaccharide biosynthesis in *Myxococcus xanthus*": All Supplementary Information

**This file contains:**

- Supplementary Figures 1-8
- Supplementary Tables 1-2
- Supplementary References

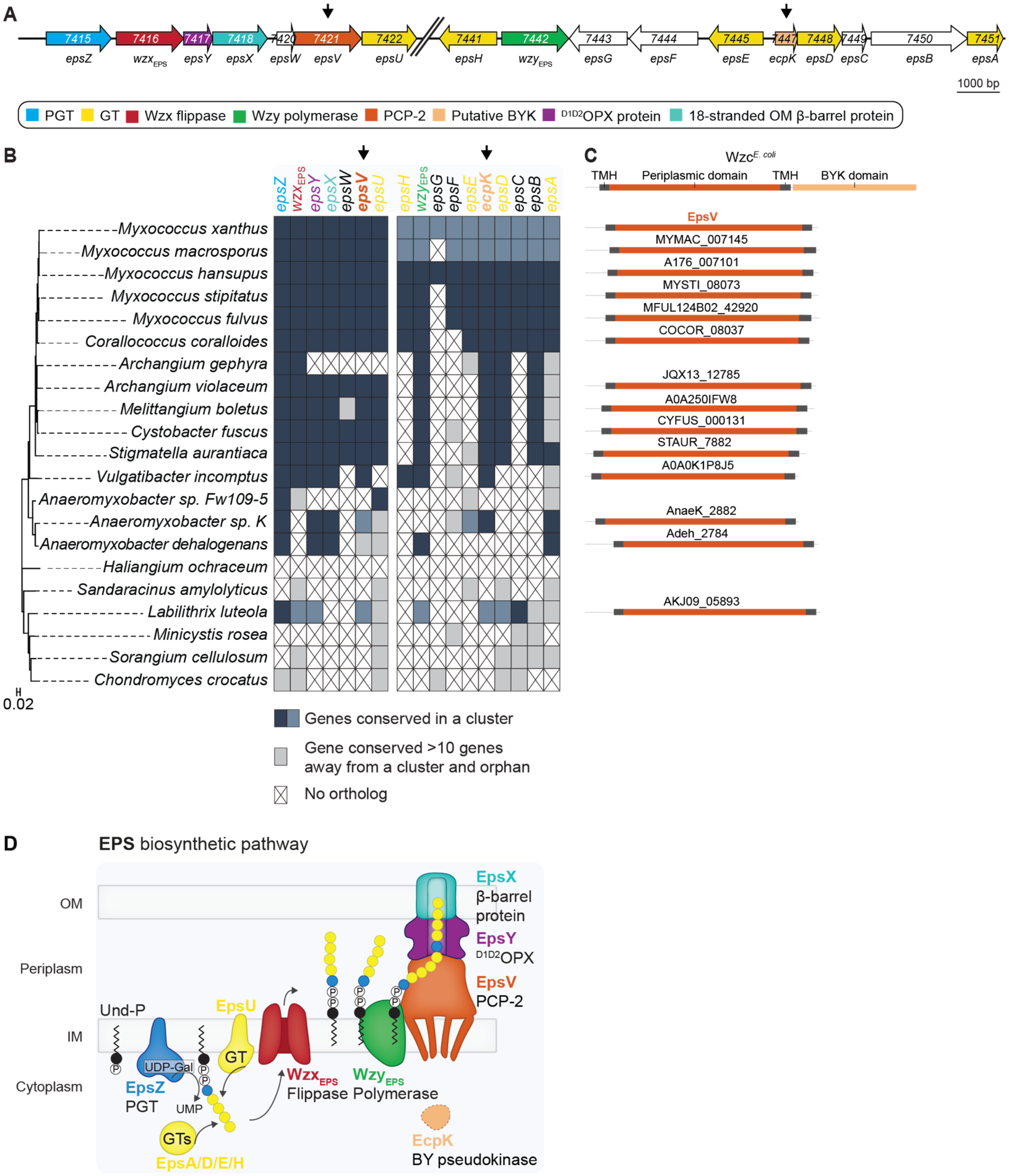


Figure S1. Myxobacterial gene clusters for EPS biosynthesis encode PCP-2 proteins lacking the BYK domain and orthologs of EcpK

(A) The two *eps* loci in *M. xanthus*. Gene names or MXAN_ numbers are indicated; genes are drawn to scale. The color code indicates the predicted functions of the gene products, shown in the lower panel, based on previous studies (1-5). EpsW is a single-domain response regulator required for EPS biosynthesis (6). *epsG* and *epsF* are annotated as encoding a magnesium transporter and a hybrid response regulator/histidine kinase, respectively. *epsF* is not important for EPS biosynthesis (5). The serine O-acetyltransferase EpsC is thought to participate in sugar nucleotide precursor biosynthesis but is not important for EPS biosynthesis (3-5). Similarly, EpsB, a predicted glycoside hydrolase, is not required for EPS biosynthesis (5).

(B) Left panel, a 16S rRNA-phylogenetic tree of fully sequenced Myxobacteria. Right panel, ortholog identification was performed using a reciprocal best hit BLASTP method as described previously (4, 7). Genes within a distance of fewer than 10 genes were considered part of the same cluster, while clusters were considered distinct when separated by more than 10 genes. Genes within the same cluster are marked with the same color. Conserved orphan genes are colored light gray, and genes without orthologs are indicated by an X. In the upper panel, *M. xanthus* genes are color-coded according to the schematic in (A). Black arrows indicate genes encoding the EpsV ortholog (orange) and the EcpK ortholog (light orange) conserved across myxobacterial *eps* gene clusters.

(C) Conservation of domain structure of PCP-2 proteins encoded in myxobacterial *eps* gene clusters. Upper panel, domain organization of the prototypical PCP-2a Wzc$^{E.\ coli}$ (8). Transmembrane helices (TMH) are shown in dark grey, the periplasmic domain in orange, and the BYK domain in light orange. Lower panel, domain structure of the PCP-2 EpsV and its myxobacterial orthologs, with domains colored according to their corresponding domains in Wzc$^{E\ coli}$.

(D) Schematic model of the EPS biosynthetic pathway in *M. xanthus*. EcpK is sometimes referred to as WzeX (2, 3, 9). Color-code as in (A). Abbreviations: UDP-Gal, uridine diphosphate galactose. UMP, uridine monophosphate.

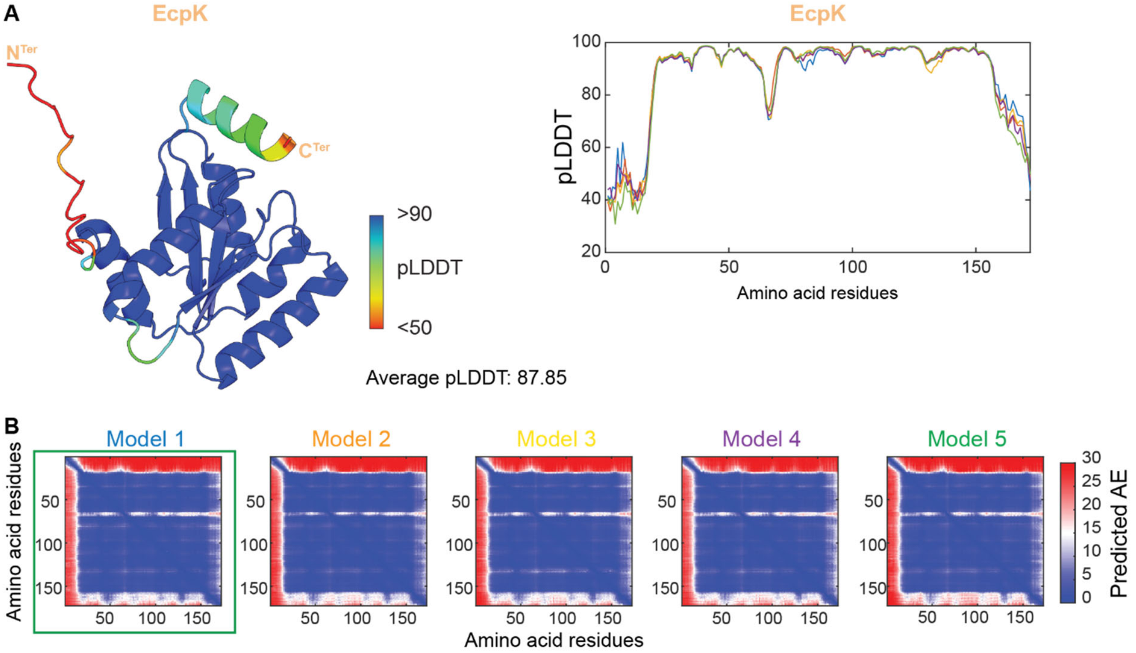


<u>Figure S2. AlphaFold2 model of EcpK.</u>

(A) Left panel, model rank 1 of EcpK colored according to pLDDT score, with average pLDDT score indicated. Right panel, pLDDT plot shown for the five generated models of EcpK.

(B) pAE plots shown for the five generated models of EcpK. Model rank 1 (highlighted by a green box) was selected for further analyses.

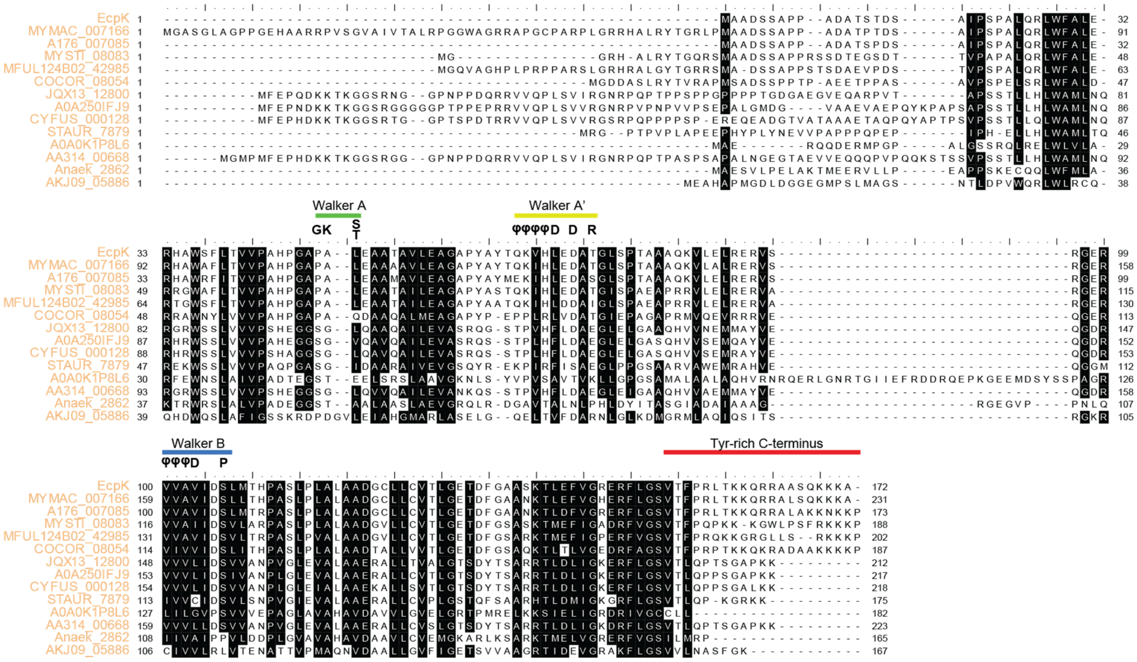


<u>Figure S3. All myxobacterial EcpK orthologs are BY pseudokinases.</u>

Sequence alignment of EcpK with its myxobacterial orthologs from Fig. S1B. The BYK signature motifs (Walker A, Walker A', Walker B) and the Tyr-rich C-terminal tail are indicated. Consensus Walker motif sequences (10) are shown, with φ representing hydrophobic residues. Amino acid residue numbering corresponds to the full-length sequences of the proteins.

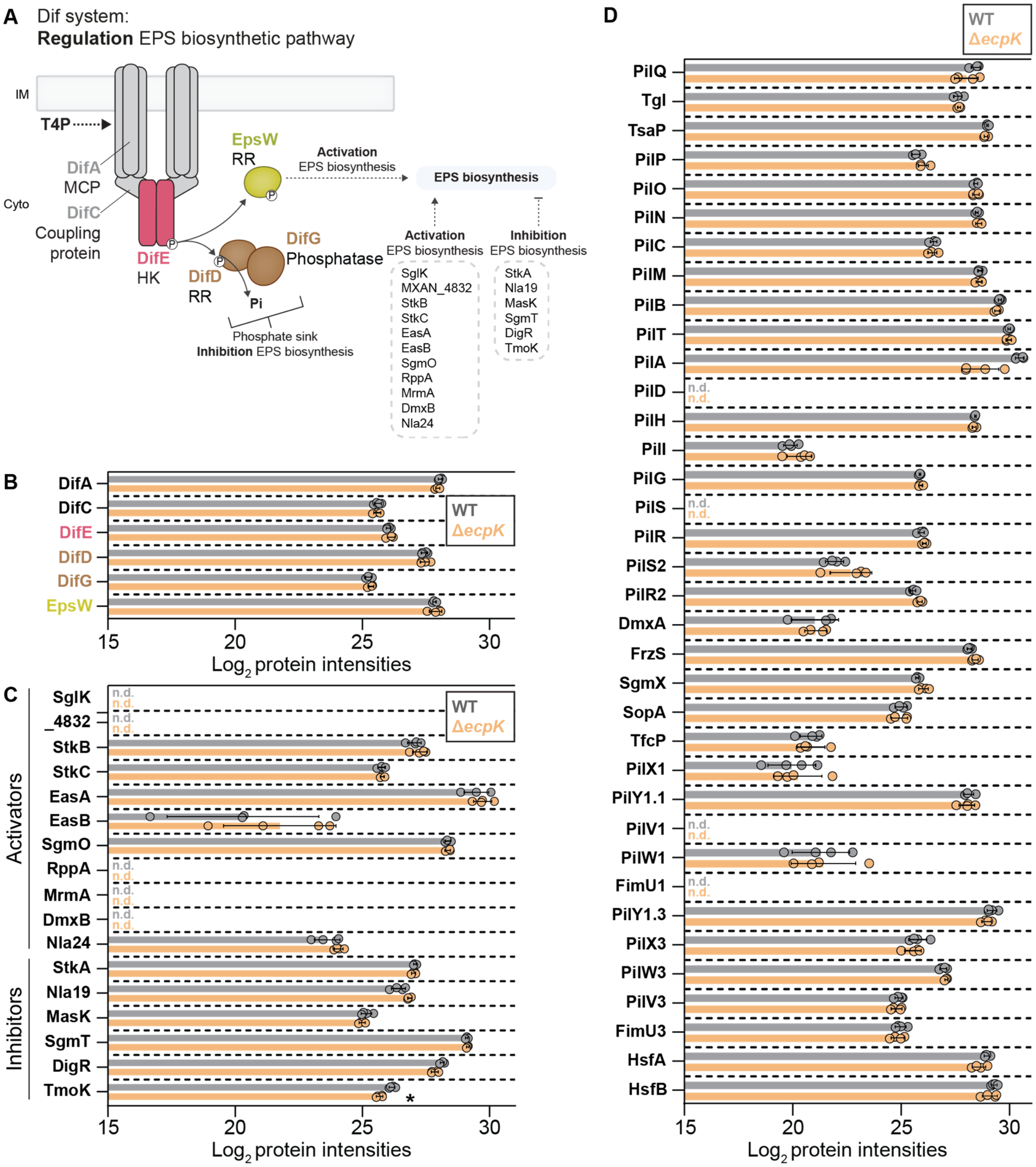


Figure S4. EcpK is not required for the accumulation of other proteins known to affect EPS biosynthesis.

(A) Schematic of the regulation of EPS biosynthesis in *M. xanthus* based on (11). Left panel, the Dif system activates EPS biosynthesis by an unknown mechanism *via* the phosphorylated response regulator EpsW, which is phosphorylated by DifE upon T4P extension (6, 12-18). DifD and DifG act as a phosphate sink, competing with EpsW for phosphorylation (16, 19). Right panel, list of additional regulators of EPS biosynthesis (11). Solid arrows indicate known mechanisms. Stippled arrows indicate unknown mechanisms. Abbreviations: MCP, methyl-accepting chemotaxis protein. HK, histidine kinase. RR, response regulator.

(B)-(D) Protein amounts in whole-cell proteomes of *M. xanthus* strains were quantified using LFQ mass spectrometry-based proteomics. Normalized $Log_2$ intensities of proteins in the indicated strains are shown. Each data point represents a biological replicate (*n* = 4 biological replicates). Error bars indicate standard deviation across these replicates. *, $p < 0.01$, Welch's test against WT. "n.d." indicates that a protein was not detected in any replicate in a strain, and is shown in the corresponding strain's color. (B) Components of the Dif system, see (A) for details. (C) Additional proteins involved in regulating EPS biosynthesis (11). (D) Proteins necessary for the assembly of a functional T4P machine. PilQ is the multimeric OM secretin stabilized by LysM-domain protein TsaP (20-23). Tgl stimulates PilQ multimerization (22, 23). PilN/-O/-P are structural components in the periplasm. PilC/-M form the IM/cytoplasmic platform complex. PilB/-T are the extension and retraction ATPases, respectively, and PilA is the major pilin (21, 24). PilH/-I/-G are suggested to form an ABC transporter (25). PilD is the prepilin leader peptidase (25, 26). PilR/-S/-R2/-S2 are regulatory proteins (27, 28). DmxA is the diguanylate cyclase important for stimulating c-di-GMP synthesis during cytokinesis and incorporation of the T4P machine at the cell pole (29). FrzS, SgmX and SopA jointly regulate T4P formation (30-35). The Cluster_1 and Cluster_3 priming complexes facilitate T4P extension (36, 37), with TfcP stabilizing PilY1.1 (38). HsfA and HsfB are a phoshphorelay involved in regulating the transcription of, among other genes (39, 40), *cluster_1* and *cluster_3* (36).

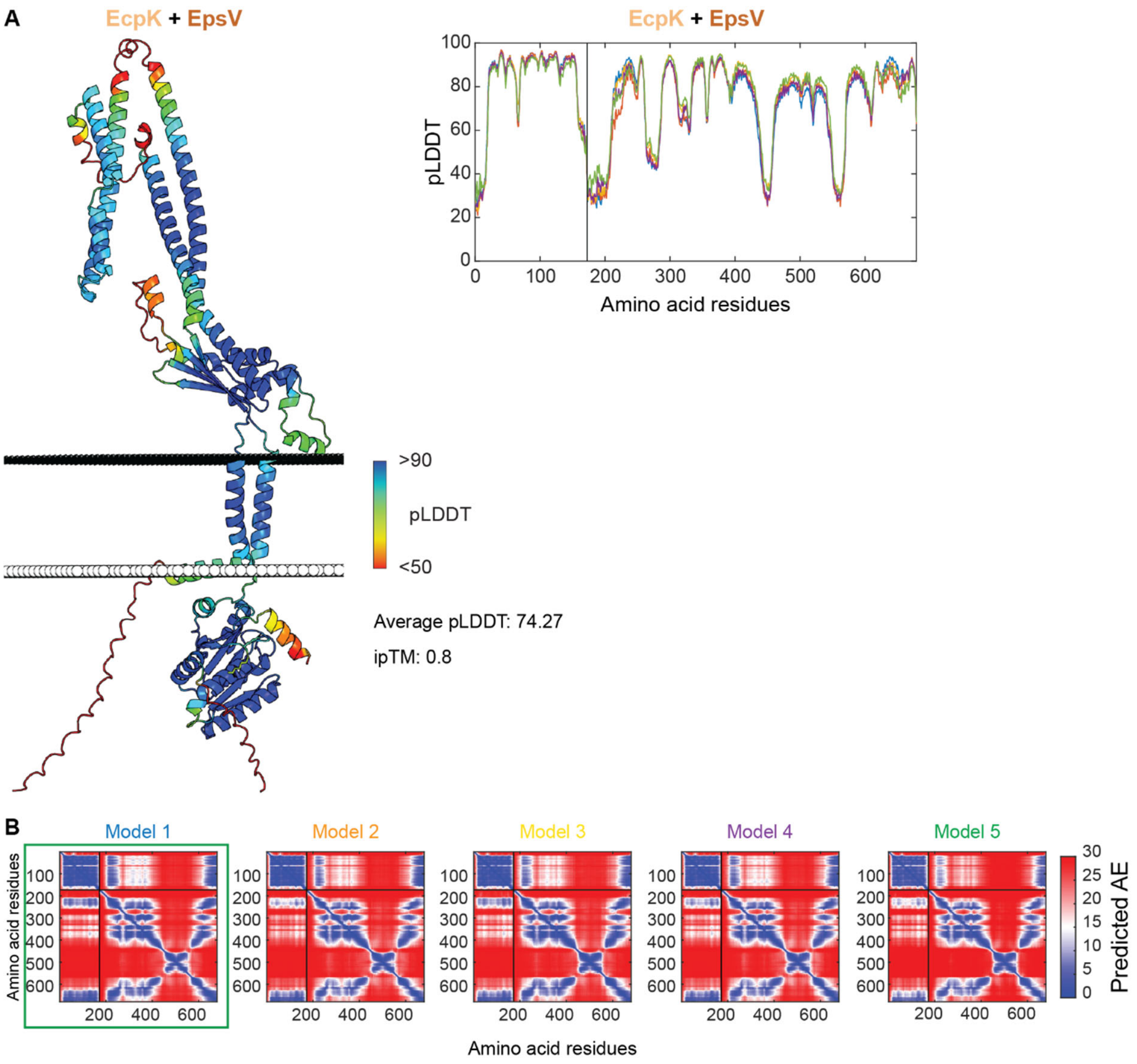


Figure S5. AlphaFold2-Multimer model of the EcpK/EpsV complex.

(A) Left panel, model rank 1 of the EcpK/EpsV complex colored according to pLDDT score, with average pLDDT and ipTM scores indicated. Right panel, pLDDT plot shown for the five generated models of the EcpK/EpsV complex.

(B) pAE plots shown for the five generated models of the EcpK/EpsV complex. Model rank 1 (highlighted by a green box) was selected for further analyses.

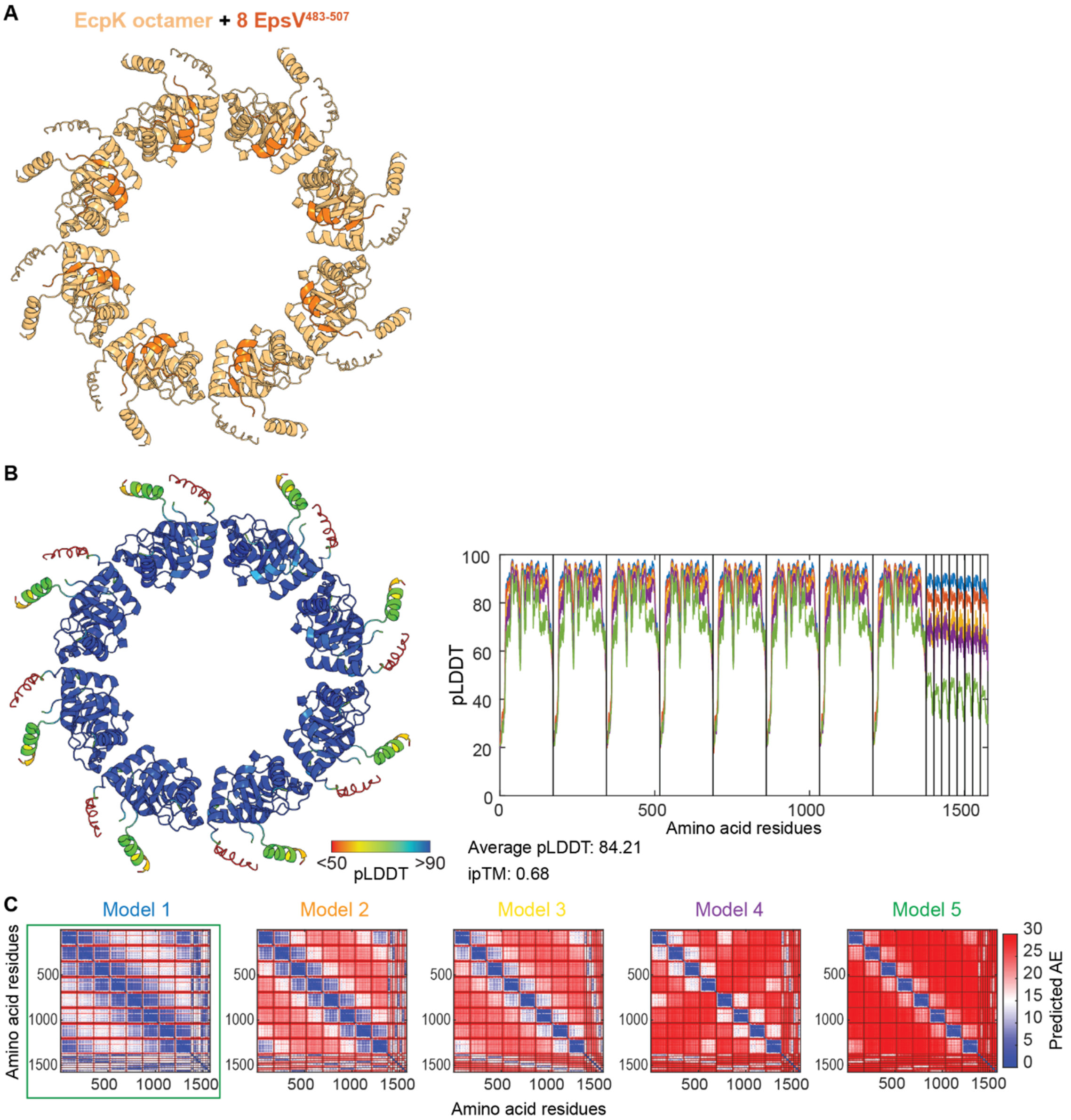


Figure S6. AlphaFold2-Multimer model of octameric EcpK/EpsV$^{483-507}$.

(A) Model rank 1 of octameric EcpK/EpsV$^{483-507}$ EcpK octamer colored in light orange, EpsV$^{483-507}$ colored in orange.

(B) Left panel, model rank 1 of octameric EcpK/EpsV$^{483-507}$ colored according to pLDDT score, with average pLDDT and ipTM scores indicated. Right panel, pLDDT plot shown for the five generated models of octameric EcpK/EpsV$^{483-507}$. Right panel, pAE plots shown for the five generated models of octameric EcpK/EpsV$^{483-507}$. Model rank 1 (highlighted by a green box) was selected for further analysis.

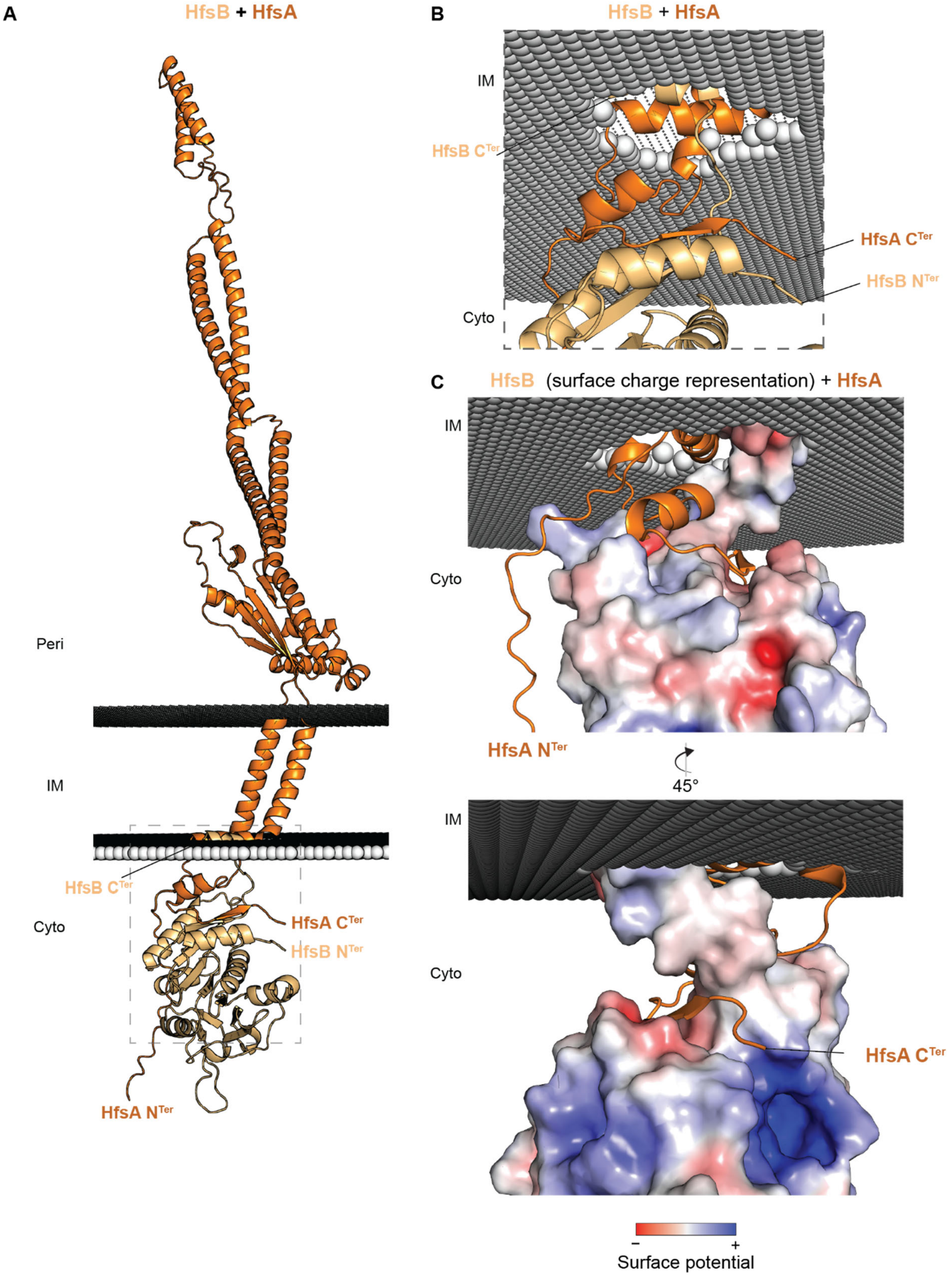


<u>Figure S7. Computational structural characterization of the HfsB-HfsA complex in *C. crescentus*.</u>

(A) AlphaFold2-Multimer model of the HfsB/HfsA complex. HfsB is shown in light orange, and HfsA in orange, with N- and C-termini highlighted. The predicted position of the HfsB/HfsA complex within the membrane was calculated using the PPM server (41). Model rank 1 is shown.

(B) Zoomed in image of the HfsB/HfsA interaction highlighting that the last 27 C-terminal, cytoplasmic residues of HfsA interacts with HfsB. Note that HfsB's C-terminal α-helix is predicted to interact with the base of one of HfsA's two TMHs. N- and C-termini of the proteins are indicated.

(C) Zoomed in image of the HfsB/HfsA interaction with HfsB shown in the surface charge representation (contoured from +5 to −5 kT $e^{-1}$), which was computed using pdb2pqr *via* the Adaptive Poisson-Boltzmann Solver server (42). Negative and positive charges are colored red and blue, respectively. Model rank 1 of the HfsA/HfsB complex, with HfsA in orange and HfsB in light-orange, respectively. N- and C-termini of HfsA are indicated.

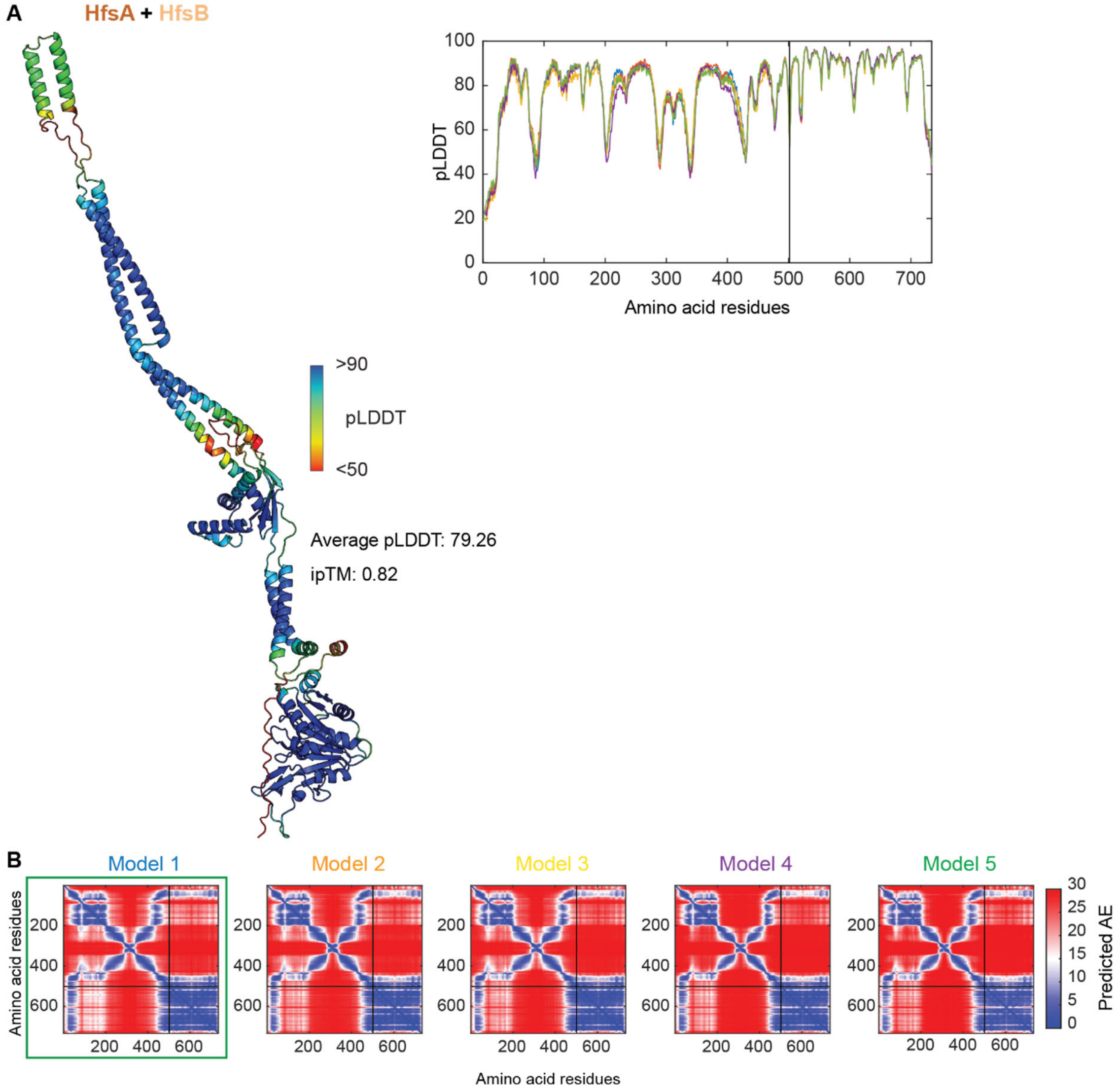


Figure S8. AlphaFold2-Multimer model of the HfsA-HfsB complex.

(A) Left panel, model rank 1 of the HfsA/HfsB complex colored according to pLDDT score, with average pLDDT and ipTM scores indicated. Right panel, pLDDT plot shown for the five generated models of the HfsA/HfsB complex. (B) pAE plots shown for the five generated models of the HfsA/HfsB complex. Model rank 1 (highlighted by a green box) was selected for further analyses.

**Table S1.** Oligonucleotides used in this work[1]

| Primer name | Sequence 5‘-3‘ | Brief description |
|---|---|---|
| LB1 | ACTGGGTACCGCTCCACGCCCATGGGTG | For Δ*ecpK* |
| LB40 | GGACGCCGCGGAATCGGCTGCCATGGG | For Δ*ecpK* |
| LB18 | AGCCGATTCCGCGGCGTCCCAGAAGAAG | For Δ*ecpK* |
| LB4 | ACTGTCTAGAGGGTCAGGGGGTTGGTCT | For Δ*ecpK* |
| LB5 | AGCACCTTGCGGTACGTG | For Δ*ecpK* |
| LB6 | GCCGCGCAGATACTCCAT | For Δ*ecpK* |
| LB7 | TGCAACGGCTGTGGTTCG | For Δ*ecpK* |
| LB8 | TGACGCTGCCGAGGAAGC | For Δ*ecpK* |
| LB9 | ACTGAAGCTTTCATGCCTTCTTCTTCTGGGACGCC | For complementation of Δ*ecpK* |
| LB10 | ACTGTCTAGAAGGCCAGGCACCGGG | For complementation of Δ*ecpK* |
| tuf2_q3-for | AGTGGAAGTCGTTGGTCTGC | Reference gene RT-qPCR |
| tuf2_q3-rev | TTGGTGTGCGGGGTGATG | Reference gene RT-qPCR |
| LB35 | AGAAGGTCCACCTGGAAGAC | RT-qPCR for *ecpK* |
| LB36 | ATCAACGAGTCGATGACGG | RT-qPCR for *ecpK* |
| 7421-q-2-for | CGACGCGGTCTTCTTTTTGA | RT-qPCR for *epsV* |
| 7421-q-2-rev | CATGATTTTGCTGACGCCCA | RT-qPCR for *epsV* |
| LB59 | ACTGAAGCTTGGCAGCCGATTCCAGCGC | BACTH with EcpK and EpsV |
| LB60 | ACTGTCTAGAGGCAGCCGATTCCAGCGC | BACTH with EcpK and EpsV |
| LB61 | ACTGAAGCTTAACGGTCCCCGCGCCCGGG | BACTH with EcpK and EpsV |
| LB62 | ACTGGGTACCGCCTTCTTCTTCTGGGAC | BACTH with EcpK and EpsV |
| LB63 | ACTGTCTAGAAACGGTCCCCGCGCCCGGGGCTC | BACTH with EcpK and EpsV |
| LB64 | ACTGGGTACCCCGCGCCGCTCCAGCTCCGC | BACTH with EcpK and EpsV |
| LB65 | ACTGAGATCTAGAGTGCGCCGCTCCAGCTCCGCCAG | BACTH with EcpK and EpsV |

[1] Underlined sequences indicate restriction sites.

**Table S2.** Fully sequenced myxobacterial genomes used for the 16S RNA tree

| Species and strain name |
|---|
| *Anaeromyxobacter dehalogenans* 2CP-C |
| *Anaeromyxobacter* sp. Fw109-5 |
| *Anaeromyxobacter* sp. K |
| *Archangium gephyra* DSM 2261 |
| *Archangium violaceum* Cb SDU34 |
| *Chondromyces crocatus* Cm c5 |
| *Corallococcus coralloides* DSM 2259 |
| *Cystobacter fuscus* DSM 52655 |
| *Haliangium ochraceum* DSM 14365 |
| *Labilithrix luteola* DSM 27648 |
| *Melittangium boletus* DSM 14713SG |
| *Minicystis rosea* DSM 24000 |
| *Myxococcus macrosporus* DSM 14675 |
| *Myxococcus hansupus* (*Myxococcus* sp. mixupus) |
| *Myxococcus stipitatus* DSM 14675 |
| *Myxococcus xanthus* DK1622 |
| *Sandaracinus amylolyticus* DSM 53668 |
| *Sorangium cellulosum* So ce 56 |
| *Stigmatella aurantiaca* DW4/3-1 |
| *Vulgatibacter incomptus* DSM 27710 |
